# Ethanol drinking sex-dependently alters cortical IL-1β synaptic signaling and cognitive behavior in mice

**DOI:** 10.1101/2024.10.08.617276

**Authors:** A Liss, MT Siddiqi, D Podder, MV Scroger, G Vessey, K Martin, NM Paperny, KT Vo, A Astefanous, N Belachew, E Idahor, FP Varodayan

## Abstract

Individuals with alcohol use disorder (AUD) struggle with inhibitory control, decision making, and emotional processing. These cognitive symptoms reduce treatment adherence, worsen clinical outcomes, and promote relapse. Neuroimmune activation is a key factor in the pathophysiology of AUD, and targeting this modulatory system is less likely to produce unwanted side effects compared to directly targeting neurotransmitter dysfunction. Notably, the cytokine interleukin-1β (IL-1β) has been broadly associated with the cognitive symptoms of AUD, though the underlying mechanisms are not well understood. Here we investigated how chronic intermittent 24-hour access two bottle choice ethanol drinking affects medial prefrontal cortex (mPFC)-related cognitive function and IL-1 synaptic signaling in male and female C57BL/6J mice. In both sexes, ethanol drinking decreased reference memory and increased mPFC IL-1 receptor 1 (IL-1R1) mRNA levels. In neurons, IL-1β can activate either pro-inflammatory or neuroprotective intracellular pathways depending on the isoform of the accessory protein (IL-1RAcP) recruited to the IL-1R1 complex. Moreover, ethanol drinking sex-dependently shifted mPFC IL-1RAcP isoform gene expression and IL-1β regulation of mPFC GABA synapses, both of which may contribute to female mPFC resiliency and male mPFC susceptibility. This type of signaling bias has become a recent focus of rational drug development. Therefore, in addition to increasing our understanding of how IL-1β sex-dependently contributes to mPFC dysfunction in AUD, our current findings also support the development of a new class of pharmacotherapeutics based on biased IL-1 signaling.

**Highlights:**

- Female mice consumed more ethanol, but were less cognitively impaired than males
- Ethanol altered IL-1β effects at mPFC GABA synapses, with females less sensitive than males
- Ethanol altered mPFC *Il1rap* mRNA to promote female neuroprotection and male neuroinflammation

## 1. Introduction

Excessive alcohol (ethanol) consumption has risen during the ongoing COVID-19 pandemic (Barbosa et al., 2021; Pollard et al., 2020; Sohi et al., 2022), and there is an urgent need to improve treatment options (Substance Abuse and Mental Health Services Administration, n.d.). Individuals with alcohol use disorder (AUD) have smaller prefrontal cortex (PFC) volumes and deficits in attention, planning, decision making, cognitive flexibility, emotional processing, inhibitory control, and learning and memory (Le Berre et al., 2017; Stavro et al., 2013). These cognitive symptoms reduce treatment adherence, worsen clinical outcomes, and promote relapse (Butler and Le Foll, 2019; Pujol et al., 2018). None of the current medications for treating AUD directly address cognitive symptoms, but there is some evidence of their positive effects on working memory, inhibitory control, and cognitive flexibility (Butler and Le Foll, 2019; Millan et al., 2012; Pujol et al., 2018). Thus, identifying novel targets that can be exploited to directly restore cognitive function represents an important avenue in AUD drug discovery.

There is growing evidence that the neuroimmune interleukin 1 (IL-1) system plays a crucial role in the cognitive symptoms associated with AUD. Human studies report functional single nucleotide polymorphisms (SNP) in IL-1 family genes that increase risk for alcohol drinking and AUD (Liu et al., 2009; Pastor et al., 2005; Saiz et al., 2009; Serretti et al., 2006). Most notably, *IL1B* C-511T (rs16944) is also associated with impaired cognition and other neuropsychiatric conditions that involve cognitive deficits (e.g., bipolar disorder, major depressive disorder, schizophrenia and dementia), and *in vitro* analyses revealed that this SNP potentiated both IL-1β gene expression and release (Chen et al., 2006; Papiol et al., 2007; Tsai, 2017; Tu et al., 2014). While IL-1β levels are low under basal conditions, immune challenges stimulate inflammasomes (innate immune system complexes) causing pro-IL-1β to be rapidly cleaved and released (Nemeth and Quan, 2021; Roberto et al., 2018). Ethanol can prime inflammasomes (Alfonso-Loeches et al., 2016, 2014; Lippai et al., 2013; Wang et al., 2015; Zou and Crews, 2012), as can IL-1β itself (Herman and Pasinetti, 2018; Patel et al., 2017), and *postmortem* studies of men with AUD report elevated brain and peripheral levels of IL-1β (Coleman et al., 2018; Crews et al., 2013; Crews and Vetreno, 2014; Vetreno et al., 2021).

Similarly, preclinical studies report increases in PFC/medial PFC (mPFC) gene and protein expression of IL-1β and its cognate receptor 1 (IL-1R1) in chronic ethanol-exposed rodents and non-human primates (Alfonso-Loeches et al., 2013; Iancu et al., 2018; Pascual et al., 2017; Pradier et al., 2018; Silva-Gotay et al., 2021; Tiwari and Chopra, 2013, 2012; Varodayan et al., 2023; Walter et al., 2020; Warden et al., 2020). The IL-1R1 complex also includes IL-1 receptor accessory protein (IL-1RAcP); upon IL-1β binding to IL-1R1, IL-1RAcP activates the myeloid differentiation primary response 88/p38 mitogen-activated protein kinase (MyD88/p38 MAPK) pathway (Bajo et al., 2019; Davis et al., 2006; Nemeth and Quan, 2021; Varodayan et al., 2023). This will trigger widespread pro-inflammatory cytokine and chemokine release, though sex differences in neuroimmune responses have been observed after alcohol intoxication in young men and women (Pascual et al., 2017), as well as in mice that drink ethanol in a moderate, binge-like manner (Alfonso-Loeches et al., 2013; Silva-Gotay et al., 2021). Interestingly, neurons express a second isoform of IL-1 accessory protein (IL-1RAcPb) that is lacking the MyD88 binding domain (Davis et al., 2006; Huang et al., 2011; Nemeth and Quan, 2021; Nguyen et al., 2011; Qian et al., 2012; Smith et al., 2009). This allows for IL-1β/IL-1R1 activation of canonical neuroprotective pathways, including phosphoinositide 3-kinase/protein kinase B (PI3K/Akt). We recently reported that heavy ethanol exposure (using 5-6 weeks of ethanol vapor) caused IL-1β to shift from a neuroprotective to proinflammatory signal in male mice (Varodayan et al., 2023), though the underlying mechanistic switch remains unknown. Here, we extend this work to examine the effects of a moderate voluntary ethanol consumption model on cognitive function and mPFC IL-1 function in both sexes and identify IL-1RAcP as a novel potential target for treating the cognitive symptoms of AUD, particularly in males.

## 2. Materials & Methods

### 2.1 Animals

We used adult male and female C57BL/6J mice (N=159; shipped at 10-12 weeks old) from The Jackson Laboratory (Bar Harbor; ME). Upon arrival, mice were given a minimum 1-week acclimation period. Mice were single-housed in a temperature- and humidity-controlled room on a 12-hour reverse light cycle (dark period was 10:00 am – 10:00 pm) for the duration of the experiment, and food and water were available *ad libitum*. All procedures were approved by Binghamton University – SUNY Institutional Animal Care and Use Committee, consistent with the National Institutes of Health Guide for the Care and Use of Laboratory Animals.

### 2.2 Chronic intermittent 24-hour access two bottle choice drinking

We used a chronic intermittent 24-hour access two bottle choice ethanol drinking (IA2BC) protocol to produce Ethanol Drinker mice that cycled between daily access to 1 bottle of ethanol and 1 bottle of water, and 2 bottles of water (Sicher et al., 2024) (**Fig. 1A**). Control mice always had 2 bottles of water. All bottles were changed in the 2-hour period spanning lights off. During the first week of exposure, the Ethanol Drinker mice received increasing ethanol concentrations (2 days of 3%, then 1 day each of 6% and 10% v/v ethanol), building up to a final concentration of 15% ethanol for the remainder of the protocol. Ethanol and water bottles were measured daily, with ethanol intake (grams of ethanol consumed in a 24-hour period divided by mouse body weight in kilograms) and ethanol preference (volume of ethanol consumed in a 24- hour period divided by total fluid consumed in the same period) were calculated. All mice were weighed weekly.

**Figure 1.**
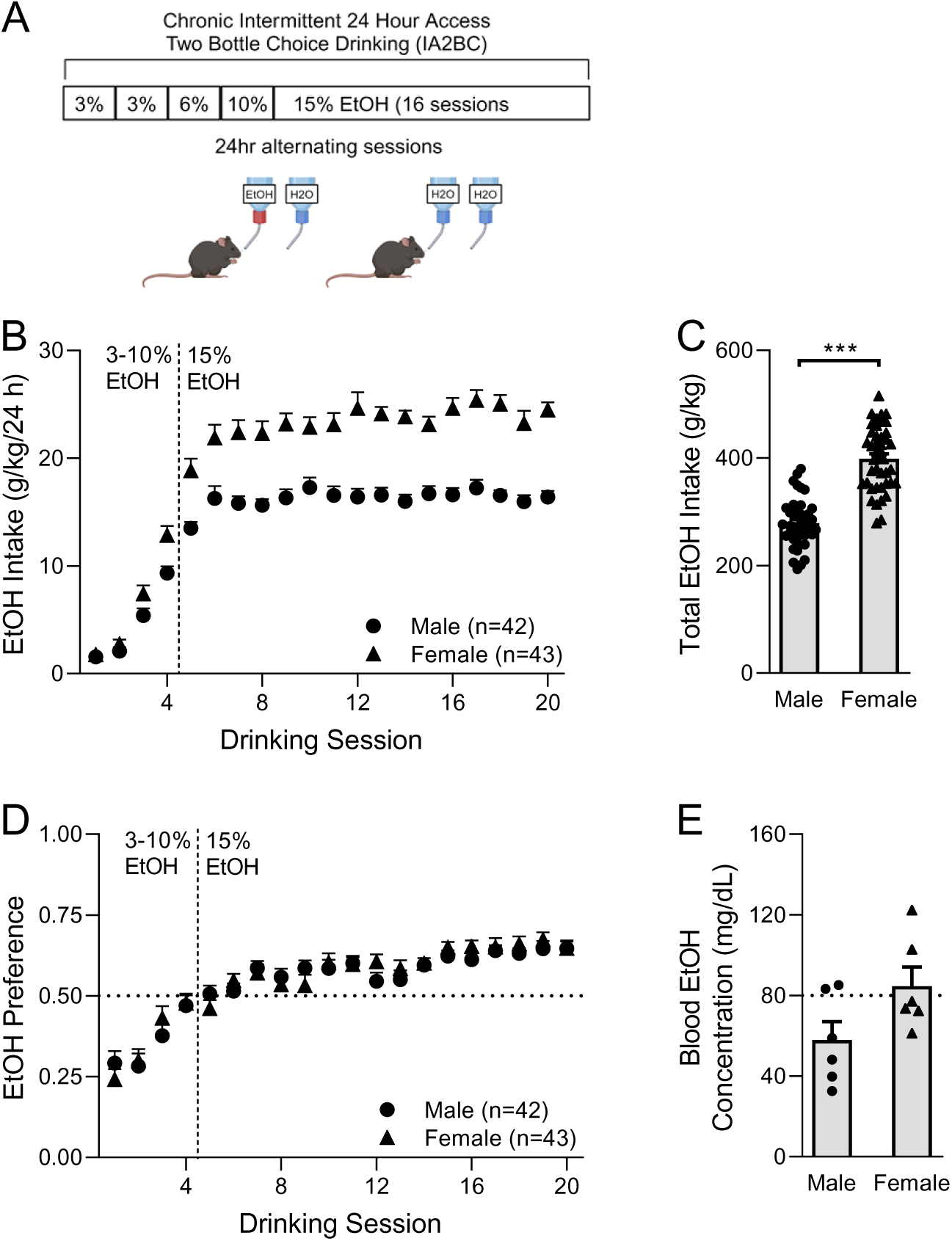
Mice consumed moderate levels of ethanol in the chronic intermittent 24-hour access two bottle choice drinking (IA2BC) protocol. A. Schematic of the IA2BC protocol. B. Daily ethanol intake escalated in both sexes. C. Female mice consumed a greater total amount of ethanol than males in the IA2BC protocol. D. Male and female mice displayed a similar increase in ethanol preference. E. BECs for 2/6 male and 5/6 female mice approached or met the criteria for binge-level drinking [∼80 mg/dL] within 4-6 hours in the final drinking session. N=42 male and 43 female mice. All data are presented as mean±SEM. \*\*\**p*<0.001 by unpaired t-test.

Mice were sacrificed 4-6 hours into their final IA2BC drinking session for blood ethanol concentration measurements (BECs), 3-5 days after IA2BC for the electrophysiology experiments, or 12 days after IA2BC for the gene expression experiments after behavioral testing. There was a main effect of sex on final body weights [sex: *F*(1, 143)=477.5, *p*<0.001; drinking: (*F*(1, 143)=0.05, *p=*0.81; interaction: *F*(1, 143)=1.45, *p=*0.23 by two-way ANOVA; (**Suppl. Fig. 1A**)]. For BECs, trunk blood was collected and centrifuged for 20 minutes at 13000 rpm at 4 °C, and the supernatants processed in a GM7 analyzer (Analox Instruments, London, UK). Four independent cohorts including both male and female mice were generated, with each cohort demonstrating similar patterns of ethanol intake and preference (**Suppl. Fig. 1B-E**).

### 2.3 Behavior

#### 2.3.1 Barnes maze

To probe the effect of IA2BC on mPFC-associated spatial reference memory, mice underwent a modified Barnes maze task with a ∼6-week gap between acquisition and retention during which IA2BC occurred (**Fig. 2A**) (Athanason et al., 2023; Gawel et al., 2019; Varodayan et al., 2018).

**Figure 2.**
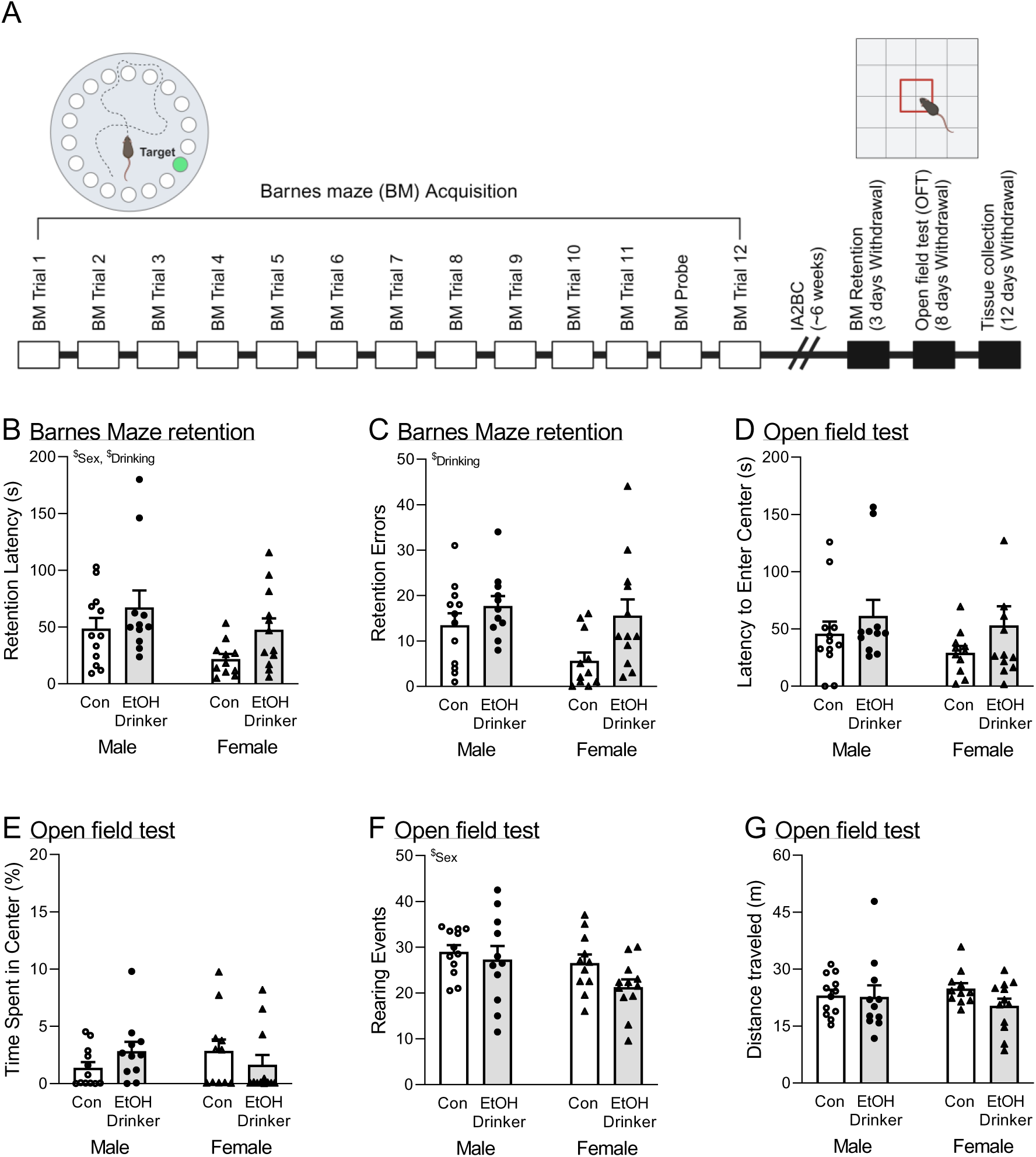
Ethanol drinking impaired cognitive performance. **A**. Schematic of the behavioral protocol used to assess reference memory (Barnes maze; BM) and anxiety-like behavior (open field test; OFT) after withdrawal from IA2BC. **B-C.** Ethanol drinking increased the (B) latency to enter the target hole and (C) number of errors made prior to entering the target hole. **D-G.** Ethanol drinking did not alter behavior in the open field test. N=11-12 mice per group. All data are presented as mean±SEM. ^$^*p*<0.05 by two- way ANOVA.

During task acquisition, mice (N=24 mice per sex) were trained to escape into a small dark recessed chamber placed under 1 of 20 small holes evenly distributed along the perimeter of an elevated, brightly lit, circular open platform (San Diego Instruments, San Diego, CA). The location of the escape box was constant for each mouse but randomized across mice, and distal visual cues aided mouse navigation. On the first day, each mouse was placed in the center of the maze, guided toward the target hole, and helped to climb in. After spending 1 min in the recessed chamber, the mouse immediately entered its first acquisition trial. For each of the 13 daily acquisition trials, each mouse was placed in the center of the maze within an opaque cylinder. After 10-20 s, the cylinder was lifted and the trial began. The trial ended once the mouse entered the escape box, and 1 min later it was returned to its home cage. If the mouse failed to enter the escape box within 3 min, it was gently guided toward the target hole and allowed to enter the escape box. Tracking videos were recorded with a ceiling-mounted video camera and analyzed with EthoVision software (Noldus, Wageningen, The Netherlands). Latency to enter the target hole and number of errors (incorrect holes visited prior to entering the target hole) were measured (**Suppl. Fig. 2A-B**).

Mice were then assigned to the Control and Ethanol Drinker groups based on their performance on the final training day. First, the mice underwent a probe test where the escape box was removed and the percentage of time each mouse spent in the target quadrant originally containing its escape box was calculated to assess spatial memory retrieval (Gawel et al., 2019; Paul et al., 2009). Each group spent >25% time in the target quadrant indicating successful task acquisition [Male Control: *t*(11)=3.41, *p*<0.01; Male EtOH Drinker: *t*(10)=3.37, *p*<0.01; Female Control: *t*(10)=4.58, *p*<0.01; Female EtOH Drinker: *t*(11)=4.53, *p*<0.001 by one sample *t*-test; (**Suppl. Fig. 2C**)], and there were no group differences [sex: *F*(1, 42)=0.49, *p*=0.49; drinking: (*F*(1, 42)=0.06, *p=*0.81; interaction: *F*(1, 42)=0.0031, *p=*0.96 by two-way ANOVA]. The mice then immediately entered their final acquisition trial, where there were no group differences in latency to enter target hole [sex: *F*(1, 42)=0.046, *p*=0.83; drinking: *F*(1, 42)=0.090, *p=*0.77; interaction: *F*(1, 42)=0.16, *p=*0.69; (**Suppl. Fig. 2D**)] or number of errors prior to target hole entry [sex: *F*(1, 42)=0.068, *p=*0.80; drinking: *F*(1, 42)=0.30, *p=*0.59; interaction: *F*(1, 42)=0.036, *p=*0.85 (**Suppl. Fig. 2E**)].

For the next ∼6 weeks, mice underwent IA2BC and did not interact with the Barnes maze apparatus. 3 full days after the last ethanol drinking session, the Barnes maze retention test was conducted under the same conditions as acquisition to assess spatial reference memory. Of note, 1 Female Control mouse and 1 Male Ethanol Drinker mouse were unexpectedly lost during IA2BC, leaving N=11-12 mice per group.

#### 2.3.2 Open field test

5 days later, the mice were tested for anxiety like behavior in the open field test (OFT) (Roberts et al., 2019). Mice were placed in the center of a brightly lit square arena and allowed to explore and allowed to explore for 5 minutes. Tracking videos were recorded with a ceiling-mounted video camera and analyzed with EthoVision software. The latency to enter the center quadrant, percentage of time spent in the center quadrant, and total distance traveled were measured. Exploratory rearing events were also scored.

### 2.4 Gene expression

The behavioral mice were sacrificed 4 days after OFT (and 12 days after their final IA2BC session) for mPFC IL-1 family gene expression analyses using real time polymerase chain reaction (rt-PCR). Briefly, mice were anesthetized using 3-5% isoflurane, and their brains were extracted, flash frozen in 2- methylbutane and stored at −80 °C (Athanason et al., 2023; Siddiqi et al., 2023). Midline mPFC micropunches (1.20 mm) were collected, and the tissue homogenized using Trizol reagent (Sigma-Aldrich, St. Louis, MO), 5 mm stainless steel beads (Qiagen, Hilden, Germany) and a TissueLyser (Qiagen). Total RNA was then extracted using RNeasy columns (Qiagen) according to manufacturer instructions, and RNA concentrations and nucleic acid purity were measured using a Nanodrop spectrophotometer (Themoscientific, Waltham, MA). cDNA was synthesized using the QuantiTect Reverse Transcription kit (Qiagen) and stored at −20 °C. Rt-PCR was performed using the cDNA template, IL-1 family gene primers (listed in **Table 1**), the CFX384 real-time PCR detection system, and iQ SYBR Green Supermix (Biorad, Hercules, CA). A single peak in the melt curve was used to ensure primer pair specificity for each gene of interest. Hypoxanthine phosphoribosyltransferase 1 (*Hprt1*) was used as a reference gene (**Suppl. Fig. 3A- B**) to normalize all gene expression data using the ΔΔC_q_ method. The Male Control group was arbitrarily designated as the ultimate control, so that the percent change could be compared across all groups. Any data points identified as outliers in the ROUT test (Q=1%) were dropped from that specific gene analysis.

**Table 1.**
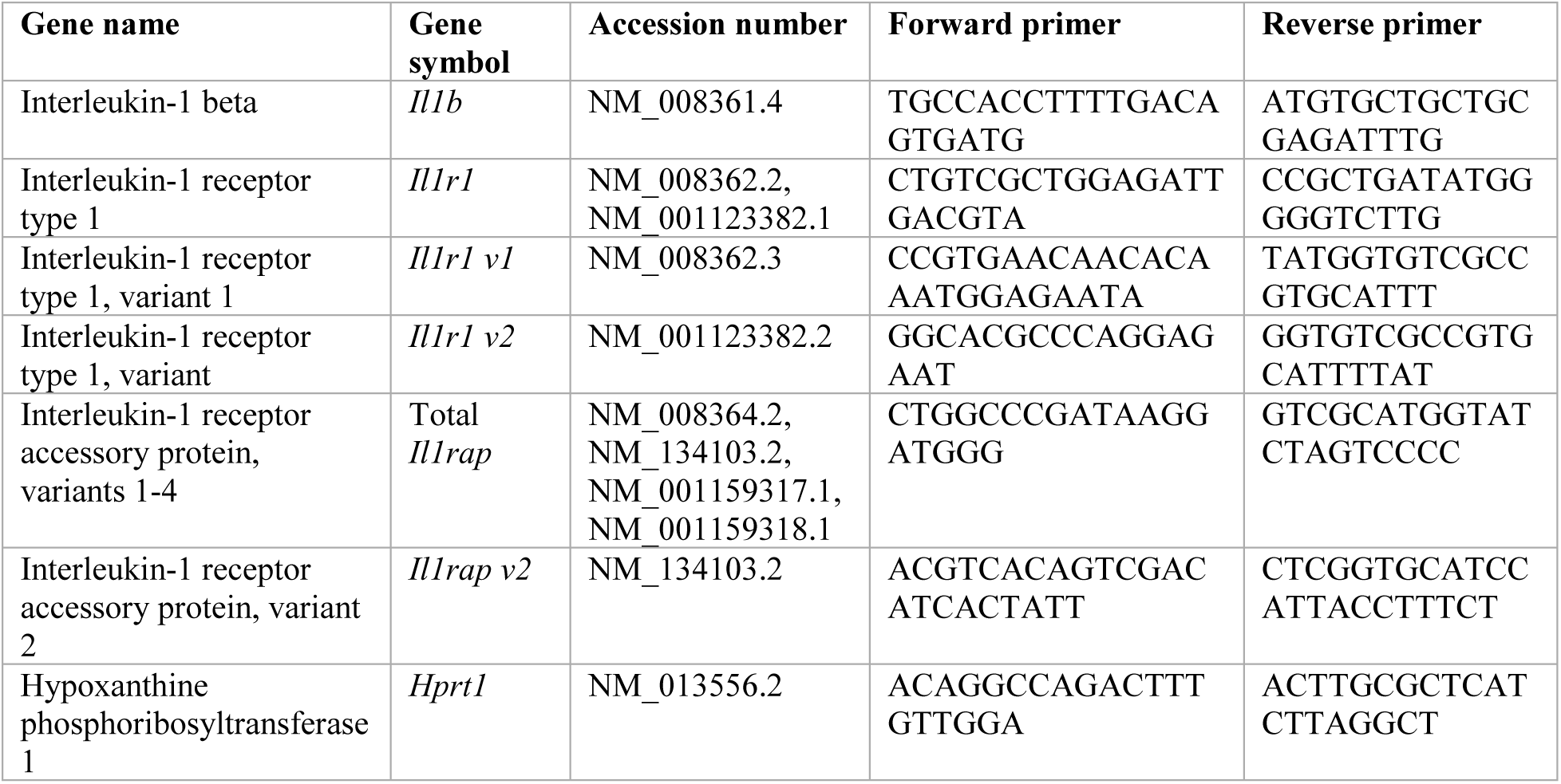
Primer pairs used for the gene expression study.

### 2.5 Electrophysiology

Mice were anesthetized with 3-5% isoflurane and decapitated (Schweitzer et al., 2016; Siddiqi et al., 2023; Varodayan et al., 2023, 2018). Extracted brains were placed in oxygenated (95% O_2_/5% CO_2_), cold high-sucrose solution (pH 7.3-7.4): 206.0 mM sucrose; 2.5 mM KCl; 0.5 mM CaCl_2_; 7.0 mM MgCl_2_; 1.2 mM NaH_2_PO_4_; 26.0 mM NaHCO_3_; 5.0 mM glucose; 5.0 mM HEPES, and sliced coronally (300-400 μm) using a Leica VT1200S vibratome (Buffalo Grove, IL). Slices were incubated in oxygenated artificial cerebrospinal fluid (aCSF): 130 mM NaCl, 3.5 mM KCl, 2 mM CaCl_2_, 1.25 mM NaH_2_PO_4_, 1.5 mM MgSO_4_, 24 mM NaHCO_3_, and 10 mM glucose for at least 1 hour at 32 °C.

In the recording chamber, each slice was perfused with 2 mL/minute room-temperature oxygenated aCSF. Prelimbic mPFC layer 2/3 neurons were visualized with infrared-differential interference contrast (IR-DIC) optics, a 40x water immersion objective (Olympus BX51WI, Tokyo, Japan) and a Retiga electro CCD camera (Teledyne, Thousand Oaks, CA). Recording neurons were located 100-300 μm from the pial surface and identified by their characteristic size and shape. Whole-cell voltage-clamp recordings from cells clamped at −70 mV were performed in gap-free acquisition mode and low-pass filtered at 10 kHz, using a Multiclamp 700B amplifier, Digidata 1550A and pClamp 10.2 software (all Molecular Devices, Sunnyvale, CA).

Pipettes (3-7 MΩ resistance) were filled with either potassium gluconate or potassium chloride internal solution: 135 mM KCl or K^+^-gluconate, 5 mM EGTA, 5 mM MgCl_2,_ 10 mM HEPES, 2 mM Mg^+^- ATP, 0.2 mM Na^+^-GTP. Spontaneous glutamatergic excitatory postsynaptic currents (sEPSCs) were pharmacologically isolated using 1 µM CGP 55845A and 30 µM bicuculline, while spontaneous γ- aminobutyric acid ionotropic receptor (GABA_A_)-mediated inhibitory postsynaptic currents (sIPSCs) were isolated with 20 µM 6,7-dinitroquinoxaline-2,3-dione (DNQX), 30 µM DL-2-amino-5-phosphonovalerate (APV) and 1 µM CGP 55845A. Series resistance was monitored with a 10 mV pulse, and recordings with a series resistance >25 MΏ or a >20% change in series resistance were excluded. Each drug was applied directly to the bath in known concentrations. We purchased APV, CGP 55845A and DNQX, from Tocris Biosciences (Ellisville, MI); recombinant mouse IL-1β from BioLegend (San Diego, CA); bicuculline from Sigma (St. Louis, MO); and ethanol from Greenfield Global (Brookfield, CT). <u>No more than one cell per mouse was used for each experimental group.</u>

sE/IPSC properties (frequency, amplitude, rise time and decay time) were analyzed during a 1.5-3 minute interval within 6-15 minutes of drug application using Mini Analysis (Synaptosoft Inc., Fort Lee, NJ) and visually confirmed. To control for cell-to-cell variation in baseline electrophysiology properties, drug effects were normalized to their own neuron’s baseline prior to group analyses. Generally, increased sE/IPSC frequencies are associated with higher release probabilities, while changes in amplitude and kinetics are linked to altered postsynaptic receptor function (Otis et al., 1994). Any cell that had a sE/IPSC property identified as an outlier in the ROUT test (Q=1%) was removed from the entire experiment. All baseline sE/IPSC properties are reported in **Suppl. Table 1.**

### 2.6 Statistical analyses

Statistical analyses were performed using one-sample and unpaired *t*-tests, Pearson correlations, and two-way ANOVAs with *post hoc* Tukey’s multiple comparisons tests where appropriate, with differences significant at p<0.05 (Prism v9.5, GraphPad, San Diego, CA). Data are represented as mean±SEM.

## 3. Results

### 3.1 Male and female mice consumed moderate levels of ethanol

We used a chronic 24-hour two bottle choice protocol to generate Ethanol Drinker and Control mice (**Fig. 1A**). While both sexes escalated their ethanol intake and preference over the 20 drinking sessions (**Fig. 1B, D**), female mice had a greater total ethanol intake [*t*(83)=10.67, *p*<0.001 by unpaired *t*-test; (**Fig. 1C**)]. We measured BECs in a subset of mice 4-6 hours into their final IA2BC ethanol drinking session and found 2/6 males and 5/6 females approached or met the criteria for binge-level drinking [∼80 mg/dL; *t*(10)=2.05, *p=*0.07; (**Fig. 1E**)].

### 3.2 Ethanol drinking impaired cognitive performance in both sexes

We have previously reported that chronic ethanol vapor exposure produced cognitive deficits in male mice (Athanason et al., 2023; Varodayan et al., 2018), so here we used a modified version of the Barnes maze task with an extended break between acquisition and retention to assess the impact of more moderate IA2BC on reference memory in both sexes (**Fig. 2A**). Male and female mice acquired the Barnes maze task similarly (see methods and **Suppl. Fig. 2**), and long-term spatial memory was assessed 3 full days after the final ethanol drinking session. We found that Ethanol Drinkers showed a retention deficit with increased latency to enter target hole [sex: *F*(1, 42)=5.13, *p*<0.05; drinking: *F*(1, 42)=4.63, *p*<0.05; interaction: *F*(1, 42)=0.12, *p=*0.73 by two-way ANOVA; (**Fig. 2B**)] and greater number of errors prior to target hole entry [sex: *F*(1, 42)=3.48, *p=*0.07; drinking: *F*(1, 42)=6.97, *p*<0.05; interaction: *F*(1, 42)=1.13, *p=*0.29 (**Fig. 2C**)], with male mice generally performing worse than females. These retention measures did not correlate with total ethanol intake in either sex.

Since the Barnes maze task is mildly aversive, five days later the mice were assessed for anxiety-like behavior in the open field test. There were no group differences in the latency to enter the center quadrant [sex: *F*(1, 42)=0.95, *p=*0.34; drinking: *F*(1, 42)=2.44, *p=*0.13; interaction: *F*(1, 42)=0.11, *p=*0.75 by two-way ANOVA; (**Fig. 2D**)] or time spent in the center quadrant [sex: *F*(1, 42)=0.35, *p=*0.56; drinking: *F*(1, 42)=0.89, *p=*0.35; interaction: *F*(1, 42)=0.89, *p=*0.35 by two-way ANOVA; (**Fig. 2E**)]. While there was a main effect of sex on exploratory rearing events [sex: *F*(1, 42)=4.21, *p<*0.05; drinking: *F*(1, 42)=2.85, *p=*0.10; interaction: *F*(1, 42)=0.73, *p=*0.40 by two-way ANOVA; (**Fig. 2F**)], total distance traveled was similar across all four groups [sex: *F*(1, 42)=0.017, *p=*0.90; drinking: *F*(1, 42)=1.45, *p=*0.24; interaction: *F*(1, 42)=1.08, *p=*0.31 by two-way ANOVA; (**Fig. 2G**)]. Overall, these data suggest that the mice may have experienced similar levels of negative affect, regardless of sex or their ethanol drinking history.

### 3.3 Ethanol drinking sex-dependently shifted mPFC IL-1RAcP isoform gene expression

Given that the interleukin 1 (IL-1) system influences PFC-associated cognitive function and its dysfunction is implicated in AUD, we next probed the impact of ethanol drinking on mPFC IL-1 family gene expression in these mice. There were no group differences in the mRNA levels of the gene encoding IL-1β (*Il1b*) [sex: *F*(1, 35)=0.003, *p=*0.96; drinking: *F*(1, 35)=0.39, *p=*0.54; interaction: *F*(1, 35)=0.12, *p=*0.73 by two-way ANOVA; (**Fig. 3A**)]. Instead, Ethanol Drinkers showed greater mPFC IL-1R1 (*Il1r1*) gene expression [sex: *F*(1, 38)=0.67, *p=*0.42; drinking: *F*(1, 38)=5.15, *p*<0.05; interaction: *F*(1, 38)=0.00001, *p=*1.00; (**Fig. 3B**)], primarily driven by the *Il1r1 variant 2* [sex: *F*(1, 40)=1.10, *p=*0.30; drinking: *F*(1, 40)=4.31, *p*<0.05; interaction: *F*(1, 40)=0.16, *p=*0.69; (**Fig. 3C**)] but not the *Il1r1 variant 1* [sex: *F*(1, 40)=7.37, *p*<0.01; drinking: *F*(1, 40)=0.69, *p=*0.16; interaction: *F*(1, 40)=0.98, *p=*0.33; (**Fig. 3D**)]. Of note, mPFC mRNA levels of *Il1r1* or either of its variants did not correlate with total ethanol intake in either sex.

**Figure 3.**
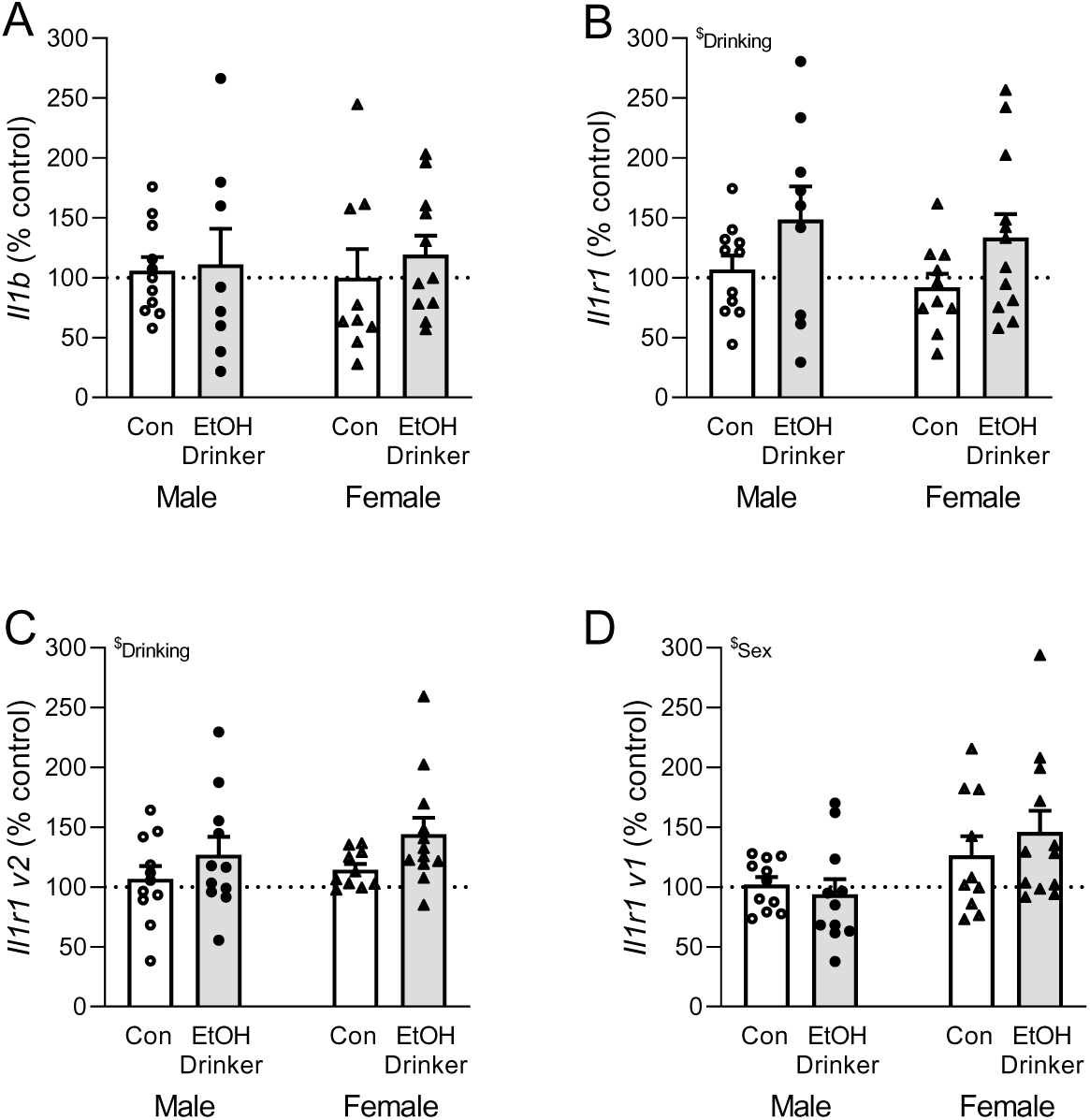
Ethanol drinking increased mPFC IL-1R1 gene expression. **A**. Ethanol drinking did not alter mPFC mRNA levels of the gene encoding IL-1β (*Il1b*). **B-D.** Ethanol drinking increased mPFC gene expression of (B) IL-1R1 (gene: *Il1r1*), primarily driven by (C) *Il1r1 variant 2* and not (D) *Il1r1 variant 1*. N=8-12 mice per group. All data are presented as mean±SEM. ^$^*p*<0.05 by two-way ANOVA.

In addition to its canonical pro-inflammatory signaling via recruitment of MyD88 by isoform a of the IL-1 receptor accessory protein (IL-RAcP; encoded by *Il1rap variant 1*), IL-1R1 can also elicit neuroprotective mechanisms via a second isoform only expressed in neurons (IL-1RAcPb; *Il1rap variant 2*) (Davis et al., 2006; Huang et al., 2011; Nemeth and Quan, 2021; Nguyen et al., 2011; Qian et al., 2012; Smith et al., 2009). mPFC transcript levels of neuroprotective *Il1rap v2* were increased in Female Ethanol Drinkers [sex: *F*(1, 40)=0.24, *p=*0.63; drinking: *F*(1, 40)=3.27, *p=*0.078; interaction: *F*(1, 40)=6.03, *p<*0.05 by two-way ANOVA; *t*(11)=2.48, *p*<0.05 by one sample *t*-test; (**Fig. 4A**)], with no group differences in total *Il1rap* [sex: *F*(1, 38)=2.29, *p=*0.14; drinking: *F*(1, 38)=0.04, *p=*0.84; interaction: *F*(1, 38)=0.04, *p=*0.84; (**Suppl. Fig. 3C**)]. When we normalized *Il1rap v2* to the total *Il1rap* mRNA levels, males had less than females [sex: *F*(1, 36)=3.58, *p*=0.067; drinking: *F*(1, 36)=5.12, *p*<0.05; interaction: *F*(1, 36)=1.09, *p=*0.30 by two-way ANOVA], though this sex difference primarily resulted from a reduction in Male Ethanol Drinkers [*t*(9)=2.50, *p*<0.05 by one sample *t*-test; (**Fig. 4B**)]. Unexpectedly, within this Male Ethanol Drinker group, normalized *Il1rap v2*/total *Il1rap* values showed a trend towards a positive correlation with Barnes maze retention errors (**Fig. 4C**).

**Figure 4.**
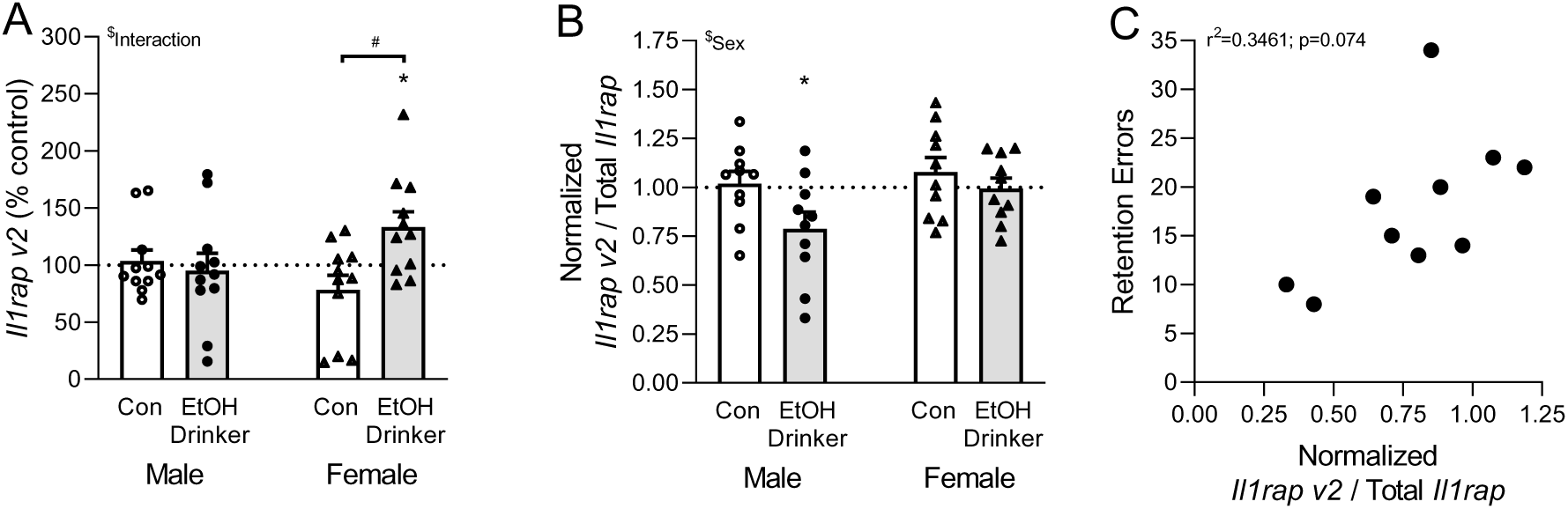
Ethanol drinking sex-dependently shifted mPFC IL-1RAcP isoform gene expression. **A**. Female Ethanol Drinkers had increased mPFC mRNA levels of the gene encoding the neuroprotective IL- 1R accessory protein (*Il1rap variant 2*). **B.** Male Ethanol Drinkers had decreased normalized *Il1rap variant 2* to the total *Il1rap* gene expression. **C.** Within the Male Ethanol Drinker group, normalized *Il1rap v2*/total *Il1rap* values showed a trend towards a positive correlation with Barnes maze retention errors. N=10-11 mice per group. All data are presented as mean±SEM. \**p*<0.05 by one-sample t-test with a hypothetical mean of 100 or 1.00. ^$^*p*<0.05 by two-way ANOVA. ^#^*p*<0.05 by Tukey’s multiple comparisons *post hoc* test.

### 3.2 IL-1β decreased mPFC inhibition in male control mice, but not females

We next studied the functional impact of IL-1 signaling on mPFC synapses. Similar to our previous work (Varodayan et al., 2023), bath application of 50 ng/mL IL-1β for 12-15 minutes decreased the sIPSC frequency onto prelimbic mPFC layer 2/3 (PL2/3) pyramidal neurons in Male Control mice [to 70.9±18.1% of baseline; *t*(10)=5.35, *p*<0.001 by one sample *t*-test; (**Fig. 5A-C**)]. There were no effects on the sIPSC amplitude or kinetics (**Fig. 5C**). Here we report for the first time that 50 ng/mL IL-1β had no effect on PL2/3 sIPSCs in the Female Control group (**Fig. 5B, D**). Using a similar experimental design, IL-1β also had no effect on PL2/3 glutamatergic sEPSCs in both sexes (**Suppl. Fig. 4**). Therefore, in basal conditions, IL-1β dampened GABA release onto PL2/3 pyramidal neurons only in males.

**Figure 5.**
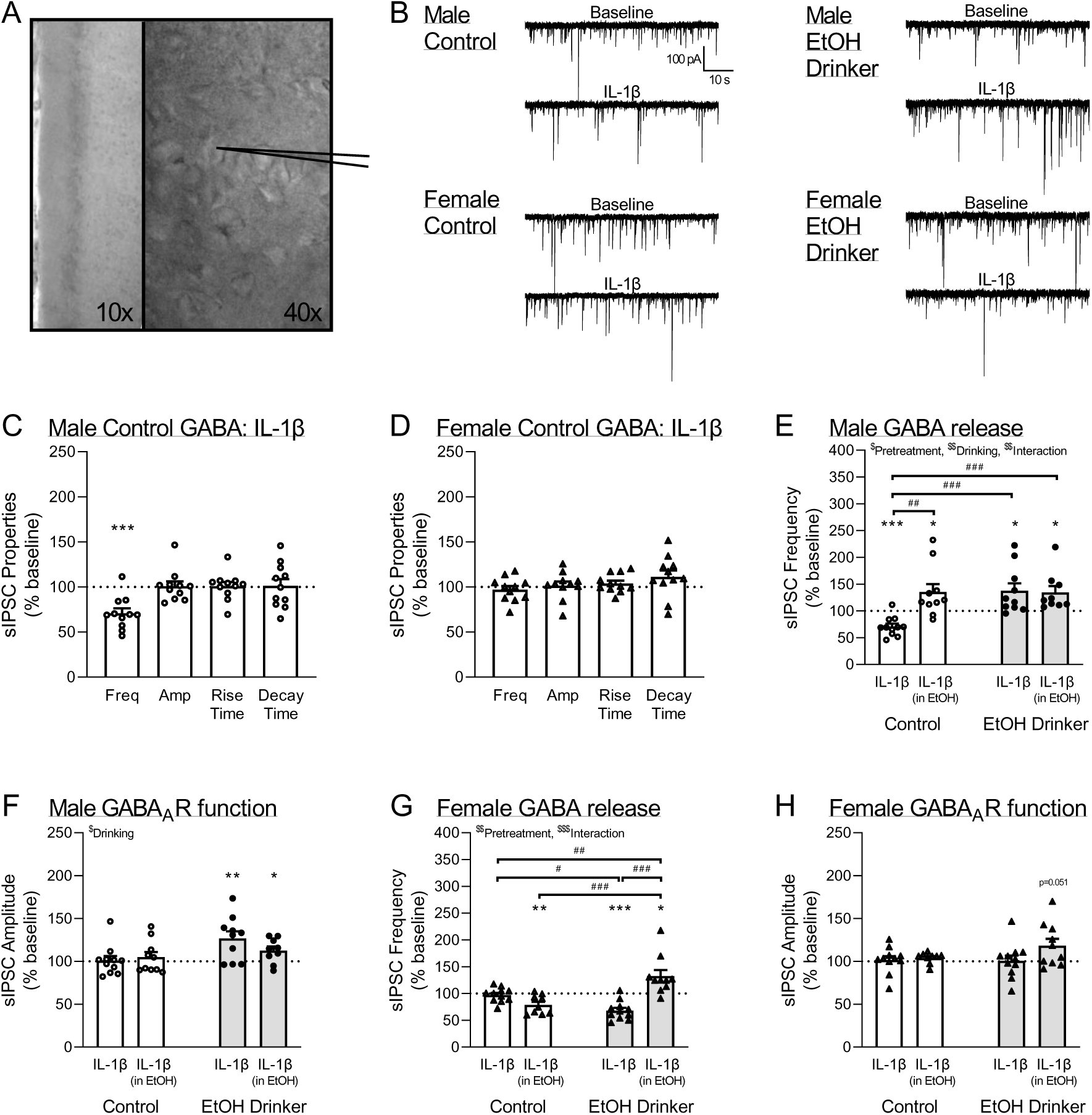
Ethanol drinking sex-dependently shifted IL-1β regulation of mPFC GABA synapses. **A.** Representative pyramidal neuron from layer 2/3 of the prelimbic mPFC (PL2/3). **B.** Representative sIPSC traces from PL2/3 pyramidal neurons before and during 50 ng/mL IL-1β application in Male and Female Control mice. **C-D**. IL-1β (C) decreased the sIPSC frequency in Male Control mice, (D) but had no effect in Female Controls. **E.** In males, IL-1β increased the PL2/3 sIPSC frequency after pretreatment with acute ethanol (in EtOH), a history of ethanol drinking mice (EtOH Drinker), or both. **F.** IL-1β increased the PL2/3 sIPSC amplitude in Male Ethanol Drinkers. **G.** In females, IL-1β decreased the PL2/3 sIPSC frequency after pretreatment with acute ethanol or a history of ethanol drinking. There was an increase in sIPSC frequency when IL-1β was applied after acute ethanol pretreatment in Female Ethanol Drinker mice. **H.** IL-1β increased the PL2/3 sIPSC amplitude after acute ethanol pretreatment in Female Ethanol Drinker mice. Note, Male and Female Control sIPSC frequency and amplitude data in panels C-D are repeated in panels E-H to allow for statistical comparisons. n=9-11 cells per group. All data are presented as mean±SEM. \**p*<0.05, \*\**p*<0.01, \*\*\**p*<0.001 by one-sample t-test with a hypothetical mean of 100. ^$^*p*<0.05, ^$$^*p*<0.01, ^$$^*p*<0.001 by two-way ANOVA. ^#^*p*<0.05, ^##^*p*<0.01, ^##^*p*<0.001 by Tukey’s multiple comparisons *post hoc* test.

### 3.3 Acute and chronic ethanol switched the IL-1β inhibitory response in male mice

As there were sex differences in IL-1β’s effects, we first opted to probe the potentially complex GABAergic interactions between ethanol and IL-1 in males. Therefore, we pretreated mPFC tissue from Male Control and 3-5 day abstinent Male Ethanol Drinkers with acute ethanol (44 mM EtOH for 10-12 minutes; (Varodayan et al., 2023)) and then co-applied IL-1β+EtOH. This experimental design allowed for direct comparison of any rapid *ex vivo* vs. persistent *in vivo* effects of ethanol on mPFC IL-1 signaling, as well as how a subsequent ethanol exposure may impact IL-1β GABAergic function after a history of drinking.

We found that ethanol altered IL-1β regulation of male PL2/3 GABA release [pretreatment: *F*(1, 36)=6.51, *p*<0.05; drinking: *F*(1, 36)=7.57, *p*<0.01; interaction: *F*(1, 36)=7.85, *p*<0.01 by two-way ANOVA and *post hoc* Tukey’s test; (**Fig. 5E**)], increasing IL-1β-induced sIPSC frequencies after acute ethanol pretreatment, chronic ethanol drinking, or both [Control/IL-1β (in EtOH): *t*(9)=2.32, *p*<0.05; EtOH Drinker/IL-1β: *t*(9)=2.70, *p*<0.05; EtOH Drinker/IL-1β (in EtOH): *t*(9)=2.88, *p*<0.05 by one sample *t*-test]. This was in contrast to the decreased sIPSC frequency we initially observed in the Male Control/IL-1β group (see section 3.2 for details). There was also a main effect of drinking on IL-1β regulation of GABA_A_ receptor function [*F*(1, 36)=7.32, *p*<0.05; (**Fig. 5F**)], with both Ethanol Drinker groups displaying increased sIPSC amplitudes [EtOH Drinker/IL-1β: *t*(9)=3.30, *p*<0.01; EtOH Drinker/IL-1β (in EtOH): *t*(9)=2.64, *p*<0.05]. Collectively, these data indicate that acute and chronic ethanol caused IL-1β to increase GABA release onto male mPFC pyramidal neurons. Ethanol drinking also enhanced IL-1β-induced postsynaptic GABA_A_ receptor function.

### 3.4 Female GABA synapses displayed a shifted sensitivity to the ethanol/IL-1β interaction

In females, a similar experimental design revealed key differences in how ethanol interacted with IL-1β regulation of GABA release [pretreatment: *F*(1, 37)=10.48, *p*<0.01; drinking: *F*(1, 37)=3.01, *p*=0.091; interaction: *F*(1, 37)=33.91, *p*<0.001; (**Fig. 5G**)]. Specifically, IL-1β decreased the sIPSC frequency in the Control/IL-1β (in EtOH) and EtOH Drinker/IL-1β groups [*t*(9)=3.60, *p*<0.01; *t*(10)=6.03, *p*<0.001, respectively], but increased it in the EtOH Drinker/IL-1β (in EtOH) group [*t*(9)=2.85, *p*<0.05]. The IL-1β-induced sIPSC amplitude of EtOH Drinker/IL-1β (in EtOH) group also approached significance, suggesting enhanced GABA_A_ receptor function [*t*(9)=2.26, *p*=0.051 (**Fig. 5H**)]. Collectively, these data indicate that either pretreatment with acute ethanol or chronic ethanol drinking (but not both) in females led to a dampening of local GABA release. Interestingly, these two effects mimicked the IL-1β response of Male Controls (see Fig. 5E, Control/IL-1β). Moreover, IL-1β’s pre- and postsynaptic effects switch after acute ethanol pretreatment in female mice with a history of chronic ethanol drinking, leading to greater overall mPFC inhibition similar to Male Ethanol Drinkers (see Fig. 5E-F, EtOH Drinker/IL-1β and EtOH Drinker/IL-1β (in EtOH)). Thus, regardless of sex, ethanol-induced IL-1 neuroadaptation fundamentally alters how the neuroimmune system shapes mPFC function, though females may be less sensitive compared to males. Future studies probing IL-1R1 intracellular signaling pathways, as well as IL-1β synaptic effects in other addiction-related brain regions, will be critical next steps towards targeted drug discovery.

## 4. Discussion

Neuroimmune activation is a key factor in the cognitive symptoms of AUD, and targeting this modulatory system is less likely to produce side effects compared to directly targeting neurotransmitter dysfunction (Crews and Vetreno, 2014; Farokhnia et al., 2019; Roberto et al., 2018). Of note, acute ethanol does not trigger a typical robust neuroimmune response. Instead, the neuroimmune system enters a primed state where inflammasomes are ready to mount exaggerated responses to future insults (Doremus-Fitzwater et al., 2015; Herman and Pasinetti, 2018; Patel et al., 2017; Siemsen et al., 2021). This heightened sensitivity is initially neuroprotective, but repeated ethanol exposure causes persistent neuroinflammation that exacerbates tissue damage and may contribute to AUD progression (Crews and Vetreno, 2014; Farokhnia et al., 2019; Roberto et al., 2018). IL-1β initiates inflammasome priming, and is also a major proinflammatory product that accumulates with priming (Herman and Pasinetti, 2018; Patel et al., 2017). *Postmortem* studies of men with AUD report elevated IL-1β and other neuroimmune signals that increase IL-1β potency (e.g. high mobility group box 1, HMGB1), some of which correlate with estimated lifetime alcohol intake (Coleman et al., 2018; Crews et al., 2013; Crews and Vetreno, 2014; Vetreno et al., 2021). This suggests that IL-1R1 may be activated with each bout of alcohol drinking and the accumulation of neuroadaptations within the IL-1 family may contribute to the impaired cognitive function that drives further drinking and AUD progression.

### 4.1 Females consumed more ethanol, but were less cognitively impaired than males

For this study we selected a chronic intermittent 24-hour access two bottle choice (IA2BC) drinking protocol in which female C57BL/6J mice had greater ethanol intake than males, though this commonly-reported sex difference (Holleran and Winder, 2017; Sicher et al., 2024) appears to be mouse strain-dependent (Bagley et al., 2021). We found that the majority of female mice and one-third of male mice approached or met the criteria for binge-level drinking; however, BECs were only collected in a subset of mice at a single timepoint and mice ethanol drinking patterns can vary based on a variety of factors including sex and basal metabolic rate (Desroches et al., 1995; Rice et al., 2023).

In both sexes, ethanol drinking led to long-term reference memory deficits in our modified Barnes maze task (∼6 week break before retention testing vs. the typical 24 hours), matching our previous results in ethanol vapor-exposed male mice (Athanason et al., 2023). There were no correlations between total ethanol intake and retention latencies or errors in either sex, suggesting that surpassing a threshold level of ethanol exposure may be enough to impair cognitive function. Of note, females, and in particular water drinking females, generally performed better relative to their male counterparts. As the Barnes maze task relies on the animal’s natural aversion to the brightly lit open space on the platform, it is possible that differences in withdrawal associated anxiety-like behavior may be a contributing factor. However, this is unlikely given our open field results where there were no group differences. Previous research has found that chronic ethanol exposure also causes mPFC-associated working memory and reversal learning deficits in both sexes (Aguirre et al., 2020; Kipp et al., 2021; Savage et al., 2000; West et al., 2018). Although the results are sometimes inconsistent, female rats showed deficits in delayed matching-to-position and males had deficits in nonmatching-to-position after chronic ethanol exposure (Savage et al., 2000). Only ethanol-exposed female rats improved their working memory performance in the delayed non-match-to-sample T-maze task (Hughes et al., 2021), and male rats out-performed females on reversal learning irrespective of ethanol exposure (Aguirre et al., 2020).

Sex differences in the cognitive sequelae of alcohol consumption have also been observed in humans, though it can be difficult to parse out the complex interaction of sex and AUD. For example, parallel studies by Sullivan, Pfefferbaum and others found that sober women with AUD had working memory deficits compared to no/low-drinking control women (Sullivan et al., 2002), while AUD produced executive dysfunction in men (Sullivan et al., 2000). It was also reported that both healthy women and women with AUD performed better than their male counterparts on tasks requiring mental flexibility, suggesting that women maintained a baseline advantage in this domain (Płotek et al., 2014; Van den Berg et al., 2017). In contrast, men and women with AUD showed similar levels of impairment in sensitivity to interference during the Stroop test (Van den Berg et al., 2017), though the results from healthy controls suggest that the deficit may actually be greater in women (Van der Elst et al., 2006). It is thought that social and biological factors that contribute to the “telescoping” effect (i.e., accelerated AUD progression) in women may underlie some of these differences (Fama et al., 2020), highlighting the need for more preclinical studies examining how different models of ethanol exposure (dosing method, frequency and pattern of exposure, duration of exposure, peak BECs, length of abstinence period, etc…) impact different cognitive domains in both sexes.

### 4.2 Female mPFC synapses were less sensitive to IL-1 regulation than males

Ethanol activates IL-1 signaling in both sexes (Crews and Vetreno, 2014; Roberto et al., 2018). Here, we found that ethanol drinking increased mPFC *Il1r1* mRNA levels (but had no effect on *Il1b*) in both sexes, which was opposite to our previous findings of increased mPFC IL-1β expression (but not IL- 1R1) in male ethanol vapor-exposed mice (Varodayan et al., 2023). These differences may stem from variation in the ethanol exposure (drinking vs. vapor, 12 days vs. 3 days withdrawal), or reflect compensatory mechanisms within the IL-1 signaling system. Also, in the present study mice underwent behavioral testing prior to sacrifice, introducing a potential confound.

Despite similar IL-1 gene expression patterns, here we found that female GABA synapses were less sensitive to IL-1β. Specifically, in basal conditions, IL-1β’s inhibitory influence was restricted to male PL2/3 synapses, where it dampened GABA release. This presynaptic effect is mediated by the neuroprotective PI3K/Akt intracellular pathway (Varodayan et al., 2023), and could potentially increase mPFC network activity. Pretreatment with bath application of acute ethanol or a history of ethanol drinking (but not both) caused female to mimic the IL-1β response of male water drinkers. This suggests the IL-1 system is already present at female PL2/3 synapses, and that ethanol can rapidly induce posttranslational changes within the IL-1R1 complex to enable its activation by IL-1β and/or its engagement of downstream PI3K/Akt signaling.

In males, ethanol rapidly and persistently switched presynaptic IL-1β function to inhibit mPFC pyramidal neurons, which could potentially disrupt top-down control. It is likely that both types of ethanol exposure (acute *ex vivo* vs. chronic *in vivo* drinking) share a common intracellular pathway since their individual effects were not additive, and our previous work implicates the MyD88/p38 MAPK pathway (Varodayan et al., 2023). In contrast, a separate IL-1 mechanism likely underlies the enhanced postsynaptic GABA_A_ receptor function observed in ethanol drinking males, even though it also contributes to greater overall inhibition of the mPFC. Notably, these same pre- and postsynaptic effects of IL-1β are observed after acute ethanol pretreatment in female mice with a history of chronic ethanol drinking, revealing an overall shifted sensitivity to IL-1 regulation of inhibitory synapses in female mice.

### 4.3 Ethanol drinking sex-dependently shifted the balance between IL-1 neuroprotective vs. pro- mechanisms

Given its key role in neuroinflammation, it is perhaps not surprising IL-1 signaling is highly regulated: 1) IL-1R2 is a decoy receptor which competitively binds IL-1β to prevent its activation of IL- 1R1, 2) IL-1RA is an endogenous competitive antagonist for IL-1β that binds IL-1R1, as well as 3) expression of neuroprotective IL-1RAcPb (encoded by *Il1rap v2)* within the IL-1R1 complex (Davis et al., 2006; Huang et al., 2011; Nemeth and Quan, 2021; Nguyen et al., 2011; Qian et al., 2012; Smith et al., 2009). We found that ethanol drinking increased mPFC *Il1rap v2* mRNA levels only in females, which might help explain their synaptic and cognitive resilience despite their greater ethanol intake. IL-1β engagement of IL-1RAcP induces proinflammatory cytokine and chemokine transcription, while IL- 1RAcPb instead regulates homeostatic processes that can be affected by sickness, including sleep and body temperature (Davis et al., 2015; J. Nguyen et al., 2019; J. T. Nguyen et al., 2019; Oles et al., 2020; Smith et al., 2009). IL-1RAcPb also curbs the canonical IL-1RAcP neuroinflammatory response by dampening its gene induction (Smith et al., 2009). After infection, IL-1RAcPb knockout mice slept less, had lower body temperature, and higher mortality compared to virus-challenged wildtype mice (Davis et al., 2015). IL-1RAcPb knockout mice were also more vulnerable to neuroimmune challenge with bacterial lipopolysaccharide, displaying greater neuroinflammatory responses and enhanced neuronal loss compared to mice deficient for both types of accessory protein or wildtype mice (Smith et al., 2009).

We also found a drop in the normalized *Il1rap v2* to the total *Il1rap* gene expression specifically in the ethanol drinking males. It is possible that this shift may explain some of the inconsistent ethanol findings observed after IL-1 pharmacological or transgenic manipulation in males. As expected, IL-1RA microinfusion into the basolateral amygdala reduced ethanol consumption in chronic exposed male mice (Marshall et al., 2016). Surprisingly, systemic IL-1RA slightly increased ethanol intake (Blednov et al., 2015), and IL-1R1 knockout and wildtype mice had similar levels of ethanol intake (Blednov et al., 2015). These disappointing results may have been driven by unintentional downregulation of IL-1RAcPb signaling. In support of this hypothesis, the normalized ratio of IL-1RAcPb to total IL-1 receptor accessory protein (i.e. IL-1RAcP + IL-1RAcPb) decreased in the aging hippocampus, leading to a stronger neuroinflammatory response and greater IL-1β-induced impairment of synaptic plasticity (Prieto et al., 2015). Systemic administration of a MyD88 mimetic that prevented its recruitment to IL-1R1 recovered preference in an object learning task in these aged mice. Since individuals with AUD show premature cortical aging (Sullivan and Pfefferbaum, 2019), an IL-1 bias towards MyD88 signaling may represent a shared substrate. Future studies should examine the impact of specifically targeting IL-1R1-induced neuroinflammatory responses on ethanol-related cognitive and other behaviors across lifespan.

Signaling bias due to conformational changes in G-protein coupled receptor complexes is well established, and has become a recent focus of rational drug development due to the potential for therapeutic benefit while preventing on-target side effects (Kenakin, 2024). Most promisingly in the addiction field, clinical studies are currently evaluating several μ-opioid receptor agonists that display bias towards G-protein vs. β-arrestin signal transduction for safer analgesic relief (Grim et al., 2020). Therefore, in addition to important implications for our understanding of how persistent IL-1β release sex-dependently contributes to mPFC dysfunction in AUD, our current findings also support the development of a new class of pharmacotherapeutics based on biased signaling within the IL-1 complex.

## Acknowledgements

This study was supported by grants from the National Institutes of Health [T32AA025606 (AL), R21AA031101 (FPV) and P50AA017823 (FPV)]; and a Health Science Transdisciplinary Award of Excellence awarded by SUNY Binghamton (FPV). The authors declare no competing financial interests.

## References

Aguirre, C.G., Stolyarova, A., Das, K., Kolli, S., Marty, V., Ray, L., Spigelman, I., Izquierdo, A., 2020. Sex-dependent effects of chronic intermittent voluntary alcohol consumption on attentional, not motivational, measures during probabilistic learning and reversal. PLoS One 15, e0234729. 10.1371/journal.pone.0234729

Alfonso-Loeches, S., Pascual, M., Guerri, C., 2013. Gender differences in alcohol-induced neurotoxicity and brain damage. Toxicology 311, 27–34. 10.1016/j.tox.2013.03.001

Alfonso-Loeches, S., Ureña-Peralta, J., Morillo-Bargues, M.J., Gómez-Pinedo, U., Guerri, C., 2016. Ethanol-Induced TLR4/NLRP3 Neuroinflammatory Response in Microglial Cells Promotes Leukocyte Infiltration Across the BBB. Neurochem Res 41, 193–209. 10.1007/s11064-015-1760-5

Alfonso-Loeches, S., Ureña-Peralta, J.R., Morillo-Bargues, M.J., Oliver-De La Cruz, J., Guerri, C., 2014. Role of mitochondria ROS generation in ethanol-induced NLRP3 inflammasome activation and cell death in astroglial cells. Front Cell Neurosci 8, 216. 10.3389/fncel.2014.00216

Athanason, A.C., Nadav, T., Cates-Gatto, C., Roberts, A.J., Roberto, M., Varodayan, F.P., 2023. Chronic ethanol alters adrenergic receptor gene expression and produces cognitive deficits in male mice. Neurobiol Stress 24, 100542. 10.1016/j.ynstr.2023.100542

Bagley, J.R., Chesler, E.J., Philip, V.M., Center for the Systems Genetics of Addiction, Jentsch, J.D., 2021. Heritability of ethanol consumption and pharmacokinetics in a genetically diverse panel of collaborative cross mouse strains and their inbred founders. Alcohol Clin Exp Res 45, 697–708. 10.1111/acer.14582

Bajo, M., Patel, R.R., Hedges, D.M., Varodayan, F.P., Vlkolinsky, R., Davis, T.D., Burkart, M.D., Blednov, Y.A., Roberto, M., 2019. Role of MyD88 in IL-1β and Ethanol Modulation of GABAergic Transmission in the Central Amygdala. Brain Sci 9, E361. 10.3390/brainsci9120361

Barbosa, C., Cowell, A.J., Dowd, W.N., 2021. Alcohol Consumption in Response to the COVID-19 Pandemic in the United States. J Addict Med 15, 341–344. 10.1097/ADM.0000000000000767

Blednov, Y.A., Benavidez, J.M., Black, M., Mayfield, J., Harris, R.A., 2015. Role of interleukin-1 receptor signaling in the behavioral effects of ethanol and benzodiazepines. Neuropharmacology 95, 309–320. 10.1016/j.neuropharm.2015.03.015

Butler, K., Le Foll, B., 2019. Impact of Substance Use Disorder Pharmacotherapy on Executive Function: A Narrative Review. Front Psychiatry 10, 98. 10.3389/fpsyt.2019.00098

Chen, H., Wilkins, L.M., Aziz, N., Cannings, C., Wyllie, D.H., Bingle, C., Rogus, J., Beck, J.D., Offenbacher, S., Cork, M.J., Rafie-Kolpin, M., Hsieh, C.-M., Kornman, K.S., Duff, G.W., 2006. Single nucleotide polymorphisms in the human interleukin-1B gene affect transcription according to haplotype context. Hum Mol Genet 15, 519–529. 10.1093/hmg/ddi469

Coleman, L.G., Zou, J., Qin, L., Crews, F.T., 2018. HMGB1/IL-1β complexes regulate neuroimmune responses in alcoholism. Brain Behav Immun 72, 61–77. 10.1016/j.bbi.2017.10.027

Crews, F.T., Qin, L., Sheedy, D., Vetreno, R.P., Zou, J., 2013. High mobility group box 1/Toll-like receptor danger signaling increases brain neuroimmune activation in alcohol dependence. Biol Psychiatry 73, 602–612. 10.1016/j.biopsych.2012.09.030

Crews, F.T., Vetreno, R.P., 2014. Neuroimmune basis of alcoholic brain damage. Int Rev Neurobiol 118, 315–357. 10.1016/B978-0-12-801284-0.00010-5

Davis, C.J., Dunbrasky, D., Oonk, M., Taishi, P., Opp, M.R., Krueger, J.M., 2015. The neuron-specific interleukin-1 receptor accessory protein is required for homeostatic sleep and sleep responses to influenza viral challenge in mice. Brain Behav Immun 47, 35–43. 10.1016/j.bbi.2014.10.013

Davis, C.N., Mann, E., Behrens, M.M., Gaidarova, S., Rebek, M., Rebek, J., Bartfai, T., 2006. MyD88- dependent and -independent signaling by IL-1 in neurons probed by bifunctional Toll/IL-1 receptor domain/BB-loop mimetics. Proceedings of the National Academy of Sciences 103, 2953–2958. 10.1073/pnas.0510802103

Desroches, D., Orevillo, C., Verina, D., 1995. Sex- and strain-related differences in first-pass alcohol metabolism in mice. Alcohol 12, 221–226. 10.1016/0741-8329(94)00098-x

Doremus-Fitzwater, T.L., Gano, A., Paniccia, J.E., Deak, T., 2015. Male adolescent rats display blunted cytokine responses in the CNS after acute ethanol or lipopolysaccharide exposure. Physiol Behav 148, 131–144. 10.1016/j.physbeh.2015.02.032

Fama, R., Le Berre, A.-P., Sullivan, E.V., 2020. Alcohol’s Unique Effects on Cognition in Women: A 2020 (Re)view to Envision Future Research and Treatment. Alcohol Res 40, 03. 10.35946/arcr.v40.2.03

Farokhnia, M., Browning, B.D., Leggio, L., 2019. Prospects for pharmacotherapies to treat alcohol use disorder: an update on recent human studies. Current Opinion in Psychiatry 32, 255–265. 10.1097/YCO.0000000000000519

Gawel, K., Gibula, E., Marszalek-Grabska, M., Filarowska, J., Kotlinska, J.H., 2019. Assessment of spatial learning and memory in the Barnes maze task in rodents-methodological consideration. Naunyn Schmiedebergs Arch Pharmacol 392, 1–18. 10.1007/s00210-018-1589-y

Grim, T.W., Acevedo-Canabal, A., Bohn, L.M., 2020. Toward Directing Opioid Receptor Signaling to Refine Opioid Therapeutics. Biol Psychiatry 87, 15–21. 10.1016/j.biopsych.2019.10.020

Herman, F.J., Pasinetti, G.M., 2018. Principles of inflammasome priming and inhibition: Implications for psychiatric disorders. Brain Behav Immun 73, 66–84. 10.1016/j.bbi.2018.06.010

Holleran, K.M., Winder, D.G., 2017. Preclinical voluntary drinking models for alcohol abstinence-induced affective disturbances in mice. Genes Brain Behav 16, 8–14. 10.1111/gbb.12338

Huang, Y., Smith, D.E., Ibanez-Sandoval, O., Sims, J.E., Friedman, W.J., 2011. Neuron-specific effects of interleukin-1β are mediated by a novel isoform of the IL-1 receptor accessory protein. Journal of Neuroscience 31, 18048–18059. 10.1523/JNEUROSCI.4067-11.2011

Hughes, B.A., O’Buckley, T.K., Boero, G., Herman, M.A., Morrow, A.L., 2021. Sex- and subtype-specific adaptations in excitatory signaling onto deep-layer prelimbic cortical pyramidal neurons after chronic alcohol exposure. Neuropsychopharmacology 46, 1927–1936. 10.1038/s41386-021-01034-1

Iancu, O.D., Colville, A., Walter, N.A.R., Darakjian, P., Oberbeck, D.L., Daunais, J.B., Zheng, C.L., Searles, R.P., McWeeney, S.K., Grant, K.A., Hitzemann, R., 2018. On the relationships in rhesus macaques between chronic ethanol consumption and the brain transcriptome. Addict Biol 23, 196–205. 10.1111/adb.12501

Kenakin, T., 2024. Bias translation: The final frontier? Br J Pharmacol 181, 1345–1360. 10.1111/bph.16335

Kipp, B.T., Nunes, P.T., Savage, L.M., 2021. Sex Differences in Cholinergic Circuits and Behavioral Disruptions Following Chronic Ethanol Exposure with and without Thiamine Deficiency. Alcohol Clin Exp Res 10.1111/acer.14594. 10.1111/acer.14594

Le Berre, A.-P., Fama, R., Sullivan, E.V., 2017. Executive Functions, Memory, and Social Cognitive Deficits and Recovery in Chronic Alcoholism: A Critical Review to Inform Future Research. Alcohol Clin Exp Res 41, 1432–1443. 10.1111/acer.13431

Lippai, D., Bala, S., Petrasek, J., Csak, T., Levin, I., Kurt-Jones, E.A., Szabo, G., 2013. Alcohol-induced IL-1β in the brain is mediated by NLRP3/ASC inflammasome activation that amplifies neuroinflammation. J Leukoc Biol 94, 171–182. 10.1189/jlb.1212659

Liu, L., Hutchinson, M.R., White, J.M., Somogyi, A.A., Coller, J.K., 2009. Association of IL-1B genetic polymorphisms with an increased risk of opioid and alcohol dependence. Pharmacogenet Genomics 19, 869–876. 10.1097/FPC.0b013e328331e68f

Marshall, S.A., Casachahua, J.D., Rinker, J.A., Blose, A.K., Lysle, D.T., Thiele, T.E., 2016. IL-1 receptor signaling in the basolateral amygdala modulates binge-like ethanol consumption in male C57BL/6J mice. Brain Behav Immun 51, 258–267. 10.1016/j.bbi.2015.09.006

Millan, M.J., Agid, Y., Brüne, M., Bullmore, E.T., Carter, C.S., Clayton, N.S., Connor, R., Davis, S., Deakin, B., DeRubeis, R.J., Dubois, B., Geyer, M.A., Goodwin, G.M., Gorwood, P., Jay, T.M., Joëls, M., Mansuy, I.M., Meyer-Lindenberg, A., Murphy, D., Rolls, E., Saletu, B., Spedding, M., Sweeney, J., Whittington, M., Young, L.J., 2012. Cognitive dysfunction in psychiatric disorders: characteristics, causes and the quest for improved therapy. Nat Rev Drug Discov 11, 141–168. 10.1038/nrd3628

Nemeth, D.P., Quan, N., 2021. Modulation of Neural Networks by Interleukin-1. Brain Plast 7, 17–32. 10.3233/BPL-200109

Nguyen, J., Gibbons, C.M., Dykstra-Aiello, C., Ellingsen, R., Koh, K.M.S., Taishi, P., Krueger, J.M., 2019. Interleukin-1 receptor accessory proteins are required for normal homeostatic responses to sleep deprivation. J Appl Physiol (1985) 127, 770–780. 10.1152/japplphysiol.00366.2019

Nguyen, J.T., Sahabandu, D., Taishi, P., Xue, M., Jewett, K., Dykstra-Aiello, C., Roy, S., Krueger, J.M., 2019. The neuron-specific interleukin-1 receptor accessory protein alters emergent network state properties in Vitro. Neurobiol Sleep Circadian Rhythms 6, 35–43. 10.1016/j.nbscr.2019.01.002

Nguyen, L., Rothwell, N.J., Pinteaux, E., Boutin, H., 2011. Contribution of interleukin-1 receptor accessory protein B to interleukin-1 actions in neuronal cells. Neurosignals 19, 222–230. 10.1159/000330803

Oles, V., Koh, K.M.S., Dykstra-Aiello, C.J., Savenkova, M., Gibbons, C.M., Nguyen, J.T., Karatsoreos, I., Panchenko, A., Krueger, J.M., 2020. Sleep- and time of day-linked RNA transcript expression in wild-type and IL1 receptor accessory protein-null mice. J Appl Physiol (1985) 128, 1506–1522. 10.1152/japplphysiol.00839.2019

Otis, T.S., De Koninck, Y., Mody, I., 1994. Lasting potentiation of inhibition is associated with an increased number of gamma-aminobutyric acid type A receptors activated during miniature inhibitory postsynaptic currents. Proc Natl Acad Sci U S A 91, 7698–7702. 10.1073/pnas.91.16.7698

Papiol, S., Molina, V., Rosa, A., Sanz, J., Palomo, T., Fañanás, L., 2007. Effect of interleukin-1beta gene functional polymorphism on dorsolateral prefrontal cortex activity in schizophrenic patients. Am J Med Genet B Neuropsychiatr Genet 144B, 1090–1093. 10.1002/ajmg.b.30542

Pascual, M., Montesinos, J., Marcos, M., Torres, J.-L., Costa-Alba, P., García-García, F., Laso, F.-J., Guerri, C., 2017. Gender differences in the inflammatory cytokine and chemokine profiles induced by binge ethanol drinking in adolescence. Addict Biol 22, 1829–1841. 10.1111/adb.12461

Pastor, I.J., Laso, F.J., Romero, A., González-Sarmiento, R., 2005. Interleukin-1 gene cluster polymorphisms and alcoholism in Spanish men. Alcohol Alcohol 40, 181–186. 10.1093/alcalc/agh153

Patel, M.N., Carroll, R.G., Galván-Peña, S., Mills, E.L., Olden, R., Triantafilou, M., Wolf, A.I., Bryant, C.E., Triantafilou, K., Masters, S.L., 2017. Inflammasome Priming in Sterile Inflammatory Disease. Trends Mol Med 23, 165–180. 10.1016/j.molmed.2016.12.007

Paul, C.-M., Magda, G., Abel, S., 2009. Spatial memory: Theoretical basis and comparative review on experimental methods in rodents. Behav Brain Res 203, 151–164. 10.1016/j.bbr.2009.05.022

Płotek, W., Łyskawa, W., Kluzik, A., Grześkowiak, M., Podlewski, R., Żaba, Z., Drobnik, L., 2014. Evaluation of the Trail Making Test and interval timing as measures of cognition in healthy adults: comparisons by age, education, and gender. Med Sci Monit 20, 173–181. 10.12659/MSM.889776

Pollard, M.S., Tucker, J.S., Green, H.D., 2020. Changes in Adult Alcohol Use and Consequences During the COVID-19 Pandemic in the US. JAMA Netw Open 3, e2022942. 10.1001/jamanetworkopen.2020.22942

Pradier, B., Erxlebe, E., Markert, A., Rácz, I., 2018. Microglial IL-1β progressively increases with duration of alcohol consumption. Naunyn Schmiedebergs Arch Pharmacol 391, 455–461. 10.1007/s00210-018-1475-7

Prieto, G.A., Snigdha, S., Baglietto-Vargas, D., Smith, E.D., Berchtold, N.C., Tong, L., Ajami, D., LaFerla, F.M., Rebek, J., Cotman, C.W., 2015. Synapse-specific IL-1 receptor subunit reconfiguration augments vulnerability to IL-1β in the aged hippocampus. Proc Natl Acad Sci U S A 112, E5078–5087. 10.1073/pnas.1514486112

Pujol, C.N., Paasche, C., Laprevote, V., Trojak, B., Vidailhet, P., Bacon, E., Lalanne, L., 2018. Cognitive effects of labeled addictolytic medications. Prog Neuropsychopharmacol Biol Psychiatry 81, 306–332. 10.1016/j.pnpbp.2017.09.008

Qian, J., Zhu, L., Li, Q., Belevych, N., Chen, Q., Zhao, F., Herness, S., Quan, N., 2012. Interleukin-1R3 mediates interleukin-1–induced potassium current increase through fast activation of Akt kinase. Proc Natl Acad Sci USA 109, 12189–12194. 10.1073/pnas.1205207109

Rice, R.C., Baratta, A.M., Farris, S.P., 2023. Home-Cage Sipper Devices Reveal Age and Sex Differences in Ethanol Consumption Patterns. bioRxiv 2023.03.22.533844. 10.1101/2023.03.22.533844

Roberto, M., Patel, R.R., Bajo, M., 2018. Ethanol and Cytokines in the Central Nervous System. Handb Exp Pharmacol 248, 397–431. 10.1007/164_2017_77

Roberts, A.J., Khom, S., Bajo, M., Vlkolinsky, R., Polis, I., Cates-Gatto, C., Roberto, M., Gruol, D.L., 2019. Increased IL-6 expression in astrocytes is associated with emotionality, alterations in central amygdala GABAergic transmission, and excitability during alcohol withdrawal. Brain Behav Immun 82, 188–202. 10.1016/j.bbi.2019.08.185

Saiz, P.A., Garcia-Portilla, M.P., Florez, G., Corcoran, P., Arango, C., Morales, B., Leza, J.C., Alvarez, S., Díaz, E.M., Alvarez, V., Coto, E., Nogueiras, L., Bobes, J., 2009. Polymorphisms of the IL-1 gene complex are associated with alcohol dependence in Spanish Caucasians: data from an association study. Alcohol Clin Exp Res 33, 2147–2153. 10.1111/j.1530-0277.2009.01058.x

Savage, L.M., Candon, P.M., Hohmann, H.L., 2000. Alcohol-induced brain pathology and behavioral dysfunction: using an animal model to examine sex differences. Alcohol Clin Exp Res 24, 465– 475.

Schweitzer, P., Cates-Gatto, C., Varodayan, F.P., Nadav, T., Roberto, M., Lasek, A.W., Roberts, A.J., 2016. Dependence-induced ethanol drinking and GABA neurotransmission are altered in Alk deficient mice. Neuropharmacology 107, 1–8. 10.1016/j.neuropharm.2016.03.003

Serretti, A., Liappas, I., Mandelli, L., Albani, D., Forloni, G., Malitas, P., Piperi, C., Zisaki, A., Tzavellas, E.O., Papadopoulou-Daifoti, Z., Prato, F., Batelli, S., Pesaresi, M., Kalofoutis, A., 2006. Interleukin-1 alpha and beta, TNF-alpha and HTTLPR gene variants study on alcohol toxicity and detoxification outcome. Neurosci Lett 406, 107–112. 10.1016/j.neulet.2006.07.003

Sicher, A.R., Liss, A., Vozella, V., Marsland, P., Seemiller, L.R., Springer, M., Starnes, W.D., Griffith, K.R., Smith, G.C., Astefanous, A., Deak, T., Roberto, M., Varodayan, F.P., Crowley, N.A., 2024. Voluntary adolescent alcohol exposure does not robustly increase adulthood consumption of alcohol in multiple mouse and rat models. bioRxiv 2024.04.30.591674. 10.1101/2024.04.30.591674

Siddiqi, M.T., Podder, D., Pahng, A.R., Athanason, A.C., Nadav, T., Cates-Gatto, C., Kreifeldt, M., Contet, C., Roberts, A.J., Edwards, S., Roberto, M., Varodayan, F.P., 2023. Prefrontal cortex glutamatergic adaptations in a mouse model of alcohol use disorder. Addict Neurosci 9, 100137. 10.1016/j.addicn.2023.100137

Siemsen, B.M., Landin, J.D., McFaddin, J.A., Hooker, K.N., Chandler, L.J., Scofield, M.D., 2021. Chronic intermittent ethanol and lipopolysaccharide exposure differentially alter Iba1-derived microglia morphology in the prelimbic cortex and nucleus accumbens core of male Long-Evans rats. J Neurosci Res 99, 1922–1939. 10.1002/jnr.24683

Silva-Gotay, A., Davis, J., Tavares, E.R., Richardson, H.N., 2021. Alcohol drinking during early adolescence activates microglial cells and increases frontolimbic Interleukin-1 beta and Toll-like receptor 4 gene expression, with heightened sensitivity in male rats compared to females. Neuropharmacology 197, 108698. 10.1016/j.neuropharm.2021.108698

Smith, D.E., Lipsky, B.P., Russell, C., Ketchem, R.R., Kirchner, J., Hensley, K., Huang, Y., Friedman, W.J., Boissonneault, V., Plante, M.-M., Rivest, S., Sims, J.E., 2009. A Central Nervous System-Restricted Isoform of the Interleukin-1 Receptor Accessory Protein Modulates Neuronal Responses to Interleukin-1. Immunity 30, 817–831. 10.1016/j.immuni.2009.03.020

Sohi, I., Chrystoja, B.R., Rehm, J., Wells, S., Monteiro, M., Ali, S., Shield, K.D., 2022. Changes in alcohol use during the COVID-19 pandemic and previous pandemics: A systematic review. Alcohol Clin Exp Res 46, 498–513. 10.1111/acer.14792

Stavro, K., Pelletier, J., Potvin, S., 2013. Widespread and sustained cognitive deficits in alcoholism: a meta-analysis. Addict Biol 18, 203–213. 10.1111/j.1369-1600.2011.00418.x

Substance Abuse and Mental Health Services Administration, n.d. Key substance use and mental health indicators in the United States: Results from the 2020 National Survey on Drug Use and Health (No. (HHS Publication No. PEP21-07-01-003, NSDUH Series H-56)). Center for Behavioral Health Statistics and Quality, Substance Abuse and Mental Health Services Administration., Rockville, MD.

Sullivan, E.V., Fama, R., Rosenbloom, M.J., Pfefferbaum, A., 2002. A profile of neuropsychological deficits in alcoholic women. Neuropsychology 16, 74–83. 10.1037//0894-4105.16.1.74

Sullivan, E.V., Pfefferbaum, A., 2019. Brain-behavior relations and effects of aging and common comorbidities in alcohol use disorder: A review. Neuropsychology 33, 760–780. 10.1037/neu0000557

Sullivan, E.V., Rosenbloom, M.J., Pfefferbaum, A., 2000. Pattern of motor and cognitive deficits in detoxified alcoholic men. Alcohol Clin Exp Res 24, 611–621.

Tiwari, V., Chopra, K., 2013. Resveratrol abrogates alcohol-induced cognitive deficits by attenuating oxidative-nitrosative stress and inflammatory cascade in the adult rat brain. Neurochem Int 62, 861–869. 10.1016/j.neuint.2013.02.012

Tiwari, V., Chopra, K., 2012. Attenuation of oxidative stress, neuroinflammation, and apoptosis by curcumin prevents cognitive deficits in rats postnatally exposed to ethanol. Psychopharmacology (Berl) 224, 519–535. 10.1007/s00213-012-2779-9

Tsai, S.-J., 2017. Effects of interleukin-1beta polymorphisms on brain function and behavior in healthy and psychiatric disease conditions. Cytokine Growth Factor Rev 37, 89–97. 10.1016/j.cytogfr.2017.06.001

Tu, P.-C., Su, T.-P., Huang, C.-C., Yang, A.C., Yeh, H.-L., Hong, C.-J., Liou, Y.-J., Liu, M.-E., Lin, C.-P., Tsai, S.- J., 2014. Interleukin-1 beta C-511T polymorphism modulates functional connectivity of anterior midcingulate cortex in non-demented elderly Han males. Brain Struct Funct 219, 61–69. 10.1007/s00429-012-0484-4

Van den Berg, J.F., Dogge, B., Kist, N., Kok, R.M., Van der Hiele, K., 2017. Gender Differences in Cognitive Functioning in Older Alcohol-Dependent Patients. Subst Use Misuse 52, 574–580. 10.1080/10826084.2016.1245341

Van der Elst, W., Van Boxtel, M.P.J., Van Breukelen, G.J.P., Jolles, J., 2006. The Stroop color-word test: influence of age, sex, and education; and normative data for a large sample across the adult age range. Assessment 13, 62–79. 10.1177/1073191105283427

Varodayan, F.P., Pahng, A.R., Davis, T.D., Gandhi, P., Bajo, M., Steinman, M.Q., Kiosses, W.B., Blednov, Y.A., Burkart, M.D., Edwards, S., Roberts, A.J., Roberto, M., 2023. Chronic ethanol induces a pro-inflammatory switch in interleukin-1β regulation of GABAergic signaling in the medial prefrontal cortex of male mice. Brain Behav Immun 110, 125–139. 10.1016/j.bbi.2023.02.020

Varodayan, F.P., Sidhu, H., Kreifeldt, M., Roberto, M., Contet, C., 2018. Morphological and functional evidence of increased excitatory signaling in the prelimbic cortex during ethanol withdrawal. Neuropharmacology 133, 470–480. 10.1016/j.neuropharm.2018.02.014

Vetreno, R.P., Qin, L., Coleman, L.G., Crews, F.T., 2021. Increased Toll-like Receptor-MyD88-NFκB- Proinflammatory neuroimmune signaling in the orbitofrontal cortex of humans with alcohol use disorder. Alcohol Clin Exp Res 45, 1747–1761. 10.1111/acer.14669

Walter, N.A.R., Zheng, C.L., Searles, R.P., McWeeney, S.K., Grant, K.A., Hitzemann, R., 2020. Chronic Voluntary Ethanol Drinking in Cynomolgus Macaques Elicits Gene Expression Changes in Prefrontal Cortical Area 46. Alcohol Clin Exp Res 44, 470–478. 10.1111/acer.14259

Wang, X., Chu, G., Yang, Z., Sun, Y., Zhou, H., Li, M., Shi, J., Tian, B., Zhang, C., Meng, X., 2015. Ethanol directly induced HMGB1 release through NOX2/NLRP1 inflammasome in neuronal cells. Toxicology 334, 104–110. 10.1016/j.tox.2015.06.006

Warden, A.S., Wolfe, S.A., Khom, S., Varodayan, F.P., Patel, R.R., Steinman, M.Q., Bajo, M., Montgomery, S.E., Vlkolinsky, R., Nadav, T., Polis, I., Roberts, A.J., Mayfield, R.D., Harris, R.A., Roberto, M., 2020. Microglia Control Escalation of Drinking in Alcohol-Dependent Mice: Genomic and Synaptic Drivers. Biological Psychiatry 88, 910–921. 10.1016/j.biopsych.2020.05.011

West, R.K., Maynard, M.E., Leasure, J.L., 2018. Binge Ethanol Effects on Prefrontal Cortex Neurons, Spatial Working Memory and Task-Induced Neuronal Activation in Male and Female Rats. Physiol Behav 188, 79–85. 10.1016/j.physbeh.2018.01.027

Zou, J., Crews, F.T., 2012. Inflammasome-IL-1β Signaling Mediates Ethanol Inhibition of Hippocampal Neurogenesis. Front Neurosci 6, 77. 10.3389/fnins.2012.00077

