## Supplemental Figures and Table for "Ethanol drinking sex-dependently alters cortical IL-1β synaptic signaling and cognitive behavior in mice"

**Affiliation:**
Developmental Exposure Alcohol Research Center and Behavioral Neuroscience Program, Department of Psychology, Binghamton University-SUNY, Binghamton, NY

**Corresponding Author:**
Dr. Florence Varodayan, Department of Psychology, Binghamton University-SUNY, S4-175G, 4400 Vestal Parkway East, Binghamton, NY 13902, USA, Tel: 607-777-4327, Fax: 607-777-4890,

### Supplemental Figures

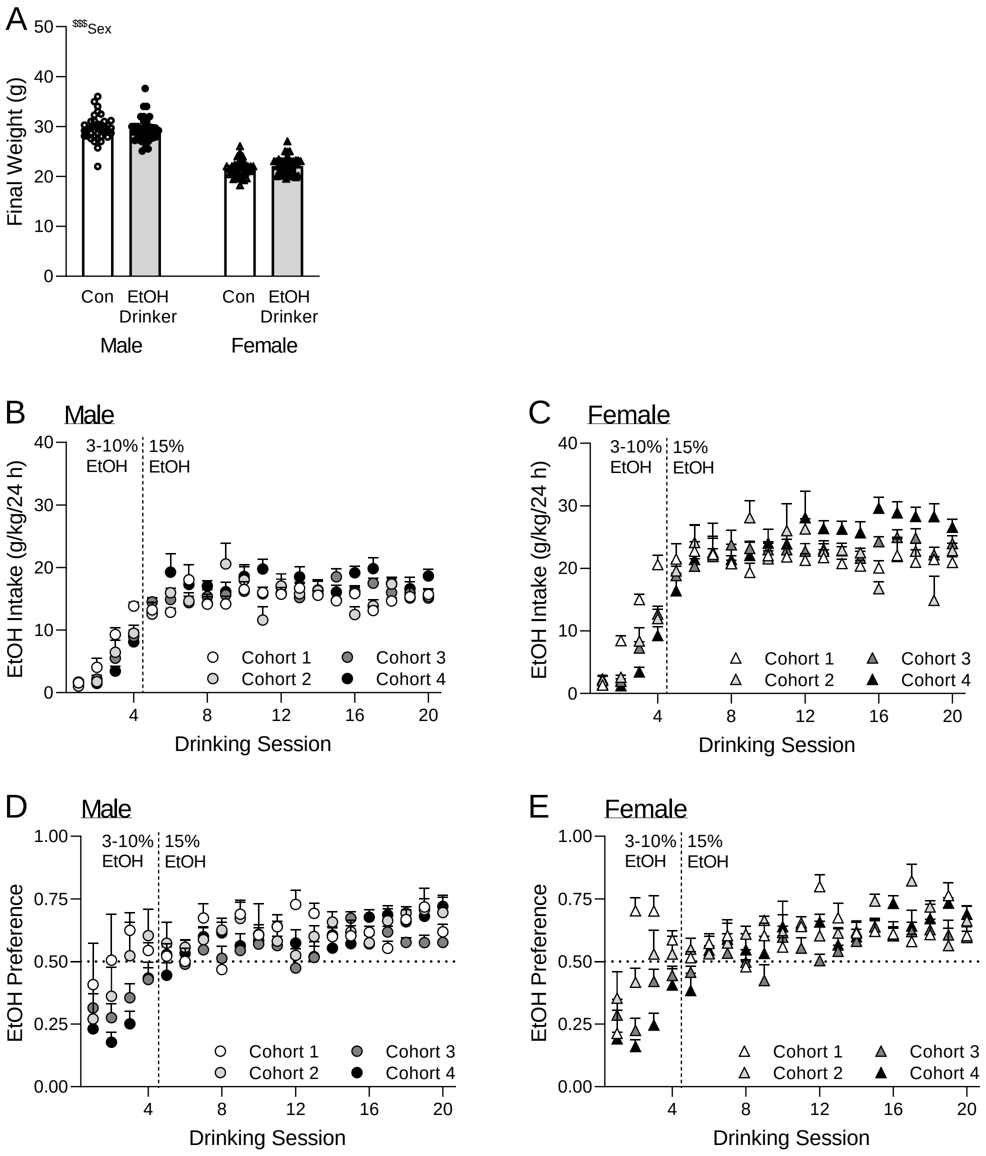


**Supplemental Figure 1. Ethanol consumption patterns in the chronic intermittent 24-hour access two bottle choice drinking (IA2BC) protocol. A.** Ethanol drinking (EtOH Drinker) did not alter the final body weights of male and female mice compared to their water drinking counterparts (Con). **B-E.** There were no cohort differences in ethanol intake or preference for either sex. Cohort 1: N=5 male and 6 female mice, cohort 2: N=6 male and 6 female mice, cohort 3: N=17 male and 16 female mice, and cohort 4: N=14 male and 15 female mice. All data are presented as mean±SEM. $^{\$\$\$}p<0.001$ by two-way ANOVA.

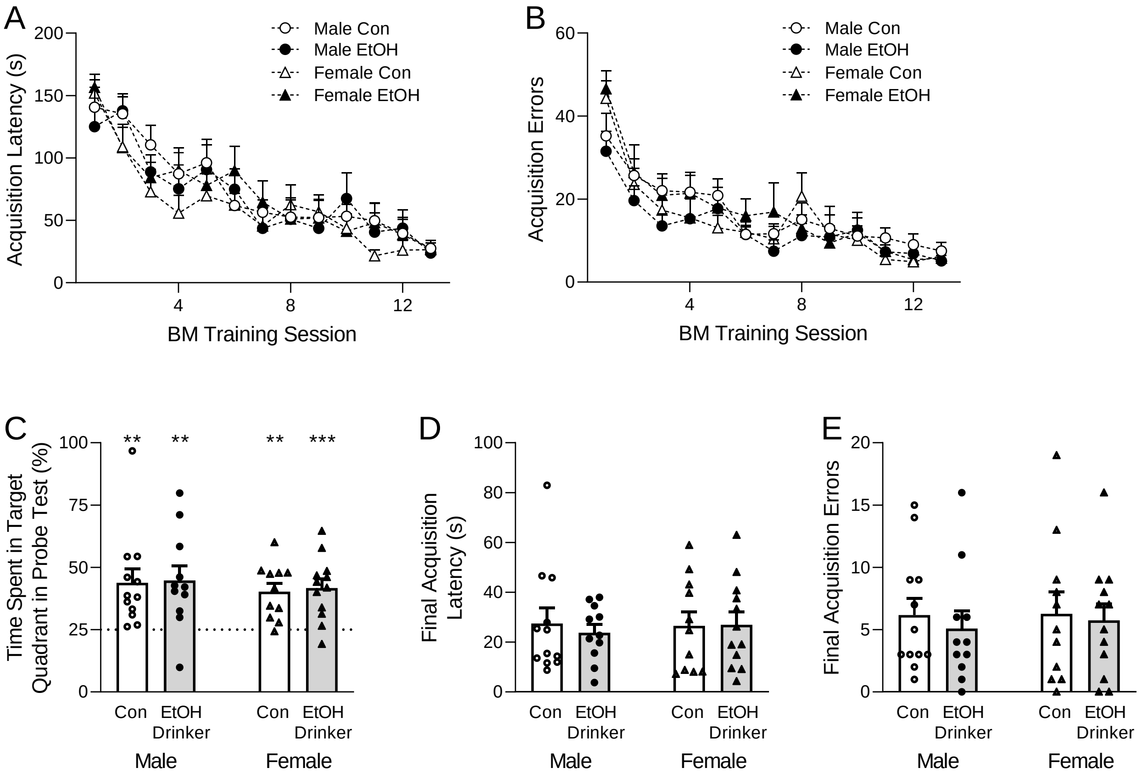


**Supplemental Figure 2. Mice were assigned to the Control and Ethanol Drinker groups based on similar Barnes maze (BM) acquisition. A-B**. Prior to ethanol exposure, all four groups showed similar BM acquisition patterns based on (B) latency to enter the target hole and (C) number of errors made prior to entering the target hole. **C.** All four groups spent more time (>25%) in the target hole quadrant vs. other quadrants during the BM probe test, indicating successful task acquisition. **D-E**. All four groups showed similar (D) latency and (E) number of errors during the final BM training session. N=11-12 mice per group. All data are presented as mean±SEM. $^{**}p<0.01$, $^{***}p<0.001$ by one-sample t-test with a hypothetical mean of 25.

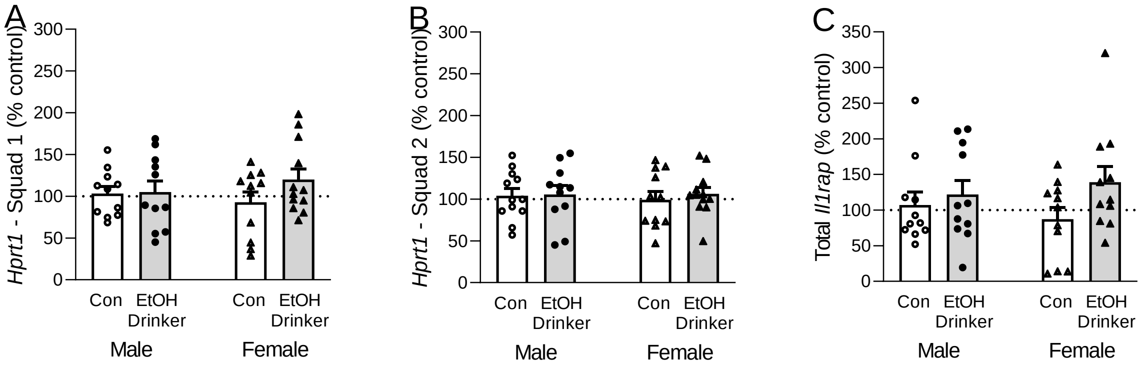


**Supplemental Figure 3. A-C.** mPFC (A-B) *Hprt1* and (C) total *Il1rap* mRNA levels were comparable across all four groups. N=11-12 mice per group.

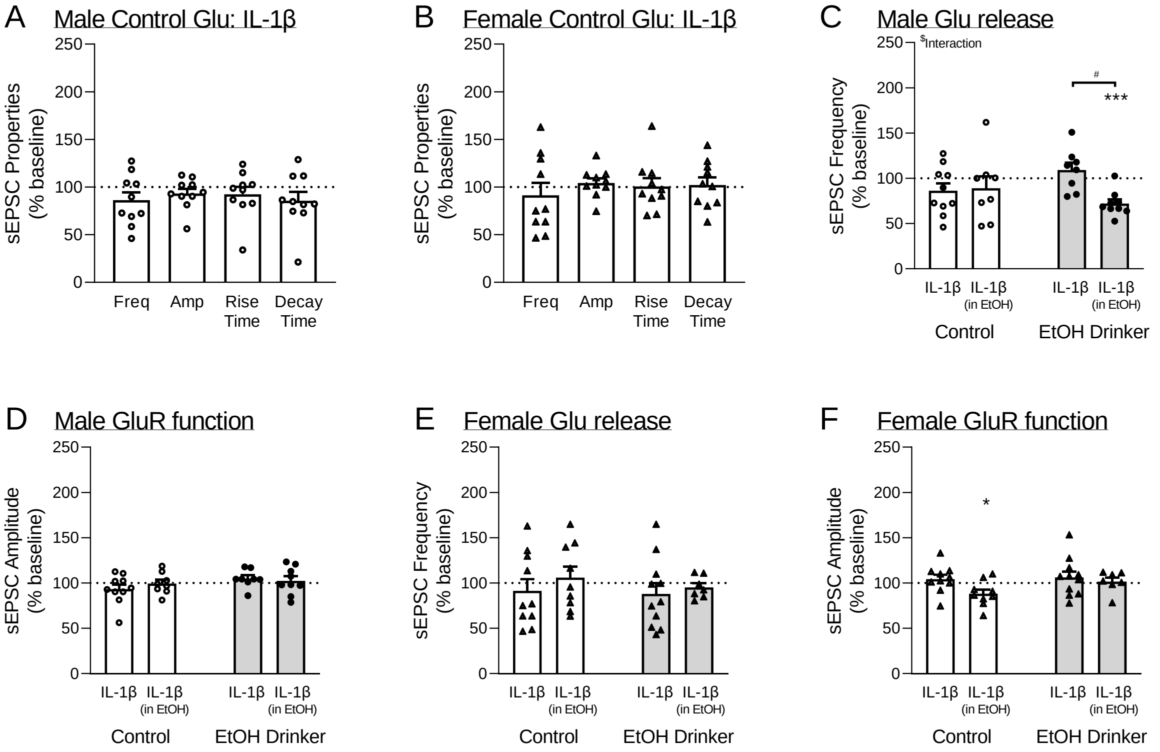


**Supplemental Figure 4. IL-1β has minimal influence at mPFC glutamate synapses. A-B**. IL-1β does not alter sEPSCs in PL2/3 pyramidal neurons in (A) Male and (B) Female Control mice. **C.** In males, there was a decrease in sEPSC frequency when IL-1β was applied after acute ethanol pretreatment in Male Ethanol Drinker mice. **D.** IL-1β does not alter sEPSC amplitudes in Male mice. **E.** IL-1β does not alter sEPSC frequencies in Female mice. **F.** IL-1β increased the PL2/3 sIPSC amplitude after acute ethanol pretreatment in Female Control mice. Note, Male and Female Control sEPSC frequency and amplitude data in panels A-B are repeated in panels C-F to allow for statistical comparisons. n=7-11 cells per group. All data are presented as mean±SEM. $^{*}p<0.05$, $^{***}p<0.001$ by one-sample t-test with a hypothetical mean of 100. $^{\$}p<0.05$ by two-way ANOVA. $^{\#}p<0.05$ by Tukey's multiple comparisons *post hoc* test.

**Supplemental Table**

| | Treatment | Frequency (Hz) | Amplitude (pA) | Rise Time (ms) | Decay Time (ms) |
|---|---|---|---|---|---|
| **sIPSCs** | **Male Control** (*n*=20 cells from 12 mice) | 3.10±0.30 | 51.5±2.4 | 2.65±0.12 | 5.66±0.35 |
| | **Male Ethanol Drinker** (*n*=20 cells from 12 mice) | 2.88±0.30 | 57.0±2.6 | 2.51±0.07 | 5.36±0.32 |
| | **Female Control** (*n*=18 cells from 13 mice) | 2.66±0.31 | 66.8±6.6$^{\$}$ | 2.48±0.11 | 6.27±0.47 |
| | **Female Ethanol Drinker** (*n*=16 cells from 13 mice) | 2.62±0.21 | 62.4±4.46$^{\$}$ | 2.57±0.08 | 6.04±0.356 |
| **sEPSCs** | **Male Control** (*n*=18 cells from 13 mice) | 0.26±0.021 | 28.5±1.0 | 1.90±0.15 | 3.17±0.25 |
| | **Male Ethanol Drinker** (*n*=20 cells from 13 mice) | 0.30±0.042 | 26.1±1.1$^{\#}$ | 1.46±0.14$^{\#}$ | 2.56±0.29 |
| | **Female Control** (*n*=17 cells from 13 mice) | 0.24±0.021 | 31.1±2.0 | 1.80±0.21 | 2.64±0.29 |
| | **Female Ethanol Drinker** (*n*=16 cells from 12 mice) | 0.22±0.028 | 26.7±1.1$^{\#}$ | 1.43±0.12$^{\#}$ | 2.59±0.27 |

**Supplemental Table 1. Basal sE/IPSC properties in prelimbic cortex layer 2/3 pyramidal neurons.** Baseline sE/IPSC frequencies, amplitudes, rise times and decay times in prelimbic layer 2/3 pyramidal neurons of Control and Ethanol Drinker mice of both sexes. There was a main effect of sex in the sIPSC amplitudes [sex: $F(1, 70)=6.01$, $p<0.05$; drinking: $F(1, 70)=0.02$, $p=0.90$; interaction: $F(1, 70)=1.40$, $p=0.24$ by two-way ANOVA], but no effects in the sIPSC frequencies [sex: $F(1, 70)=1.50$, $p=0.22$; drinking: $F(1, 70)=0.20$, $p=0.66$; interaction: $F(1, 70)=0.10$, $p=0.76$ by two-way ANOVA], rise times [sex: $F(1, 70)=0.29$, $p=0.59$; drinking: $F(1, 70)=0.07$, $p=0.79$; interaction: $F(1, 70)=1.47$, $p=0.23$ by two-way ANOVA], or decay times [sex: $F(1, 70)=2.87$, $p=0.09$; drinking: $F(1, 70)=0.50$, $p=0.48$; interaction: $F(1, 70)=0.01$, $p=0.92$ by two-way ANOVA]. There were main effects of ethanol drinking in the sEPSC amplitudes [sex: $F(1, 67)=1.35$, $p=0.25$; drinking: $F(1, 67)=6.59$, $p<0.05$; interaction: $F(1, 67)=0.56$, $p=0.46$ by two-way ANOVA] and rise times [sex: $F(1, 67)=0.20$, $p=0.69$; drinking: $F(1, 67)=6.57$, $p<0.05$; interaction: $F(1, 67)=0.05$, $p=0.82$ by two-way ANOVA], but no effects in the sIPSC frequencies [sex: $F(1, 67)=2.75$, $p=0.10$; drinking: $F(1, 67)=0.03$, $p=0.86$; interaction: $F(1, 67)=0.83$, $p=0.37$ by two-way ANOVA], or decay times [sex: $F(1, 67)=0.83$, $p=0.36$; drinking: $F(1, 67)=1.42$, $p=0.24$; interaction: $F(1, 67)=0.99$, $p=0.32$ by two-way ANOVA]. All data are presented as mean±SEM. $^{\$}p<0.05$ main effect of sex, $^{\#}p<0.05$ main effect of treatment by two-way ANOVA.
